# Dysregulation of the *de novo* proteome accompanies pathology progression in the APP/PS1 mouse model of Alzheimer’s disease

**DOI:** 10.1101/2020.08.21.260786

**Authors:** Megan K. Elder, Hediye Erdjument-Bromage, Mauricio M. Oliveira, Maggie Mamcarz, Thomas A. Neubert, Eric Klann

## Abstract

Alzheimer’s disease (AD) is an age-related neurodegenerative disorder, but neuropathological changes in AD begin years before memory impairment. Investigation of the early molecular abnormalities in AD might offer innovative opportunities to target memory impairment prior to its onset. Decreased protein synthesis plays a fundamental role in AD, yet the consequences for cellular function remain unknown. We hypothesize that alterations in the *de novo* proteome drive early metabolic changes in the hippocampus that persist throughout AD progression. Using a combinatorial amino acid tagging approach to selectively label newly synthesized proteins, we found that the *de novo* proteome is disturbed in APP/PS1 mice prior to pathology development, affecting the synthesis of multiple components of synaptic, lysosomal and mitochondrial pathways. Furthermore, the synthesis of large clusters of ribosomal subunits were affected throughout neuropathology development in these mice. Our data suggest that large-scale changes in protein synthesis could underlie cellular dysfunction in AD.

## Introduction

Alzheimer’s disease (AD) is an age-related neurodegenerative condition characterized by progressive and devastating cognitive impairment. AD is classically characterized by extensive deposition of amyloid beta (Aβ) plaques and intracellular inclusions of hyperphosphorylated tau, in the form of neurofibrillary tangles (Alzheimer, 1907). In addition to the deposition of plaques and tangles, AD is characterized by extensive atrophy of the brain, which follows a prescribed trajectory throughout disease progression. Initially atrophy is restricted to the hippocampi and medial temporal lobes, even before the appearance of symptoms (Scahill et al., 2002). The cause of this neuronal atrophy has not yet been elucidated, but it is likely that pathological alterations in protein synthesis are a contributing factor (Hernández-Ortega et al., 2016). In a healthy biological system, proteins are made and degraded in a controlled process termed protein homeostasis (proteostasis). However, the intrinsic and extrinsic stressors that accumulate with age and disease can challenge the delicate balance between protein synthesis and degradation (Kaushik and Cuervo, 2015).

Removal of misfolded or otherwise damaged proteins is essential to avoid toxicity and is regulated by a variety of proteolytic systems, whose dysfunction has been implicated in AD pathophysiology (Ciechanover and Kwon, 2015). Proteasomal activity is perturbed in vulnerable cortical areas and the hippocampus in late stage AD (Keller et al., 2000), whereas proteolysis has been shown to be impaired in very mild stages of the disease, suggesting that proteostasis disruption might occur prior to the development of symptoms (Mawuenyega et al., 2010).

On the opposite side of the proteostatic balance is protein synthesis. Following transcription, mRNA is translated to a polypeptide chain via ribosomal activity, a process that in neurons occurs both somatically and locally in dendrites and axons (Holt et al., 2019). This *de novo* protein synthesis is a dynamic process, as it is necessary for basal neuronal function, responding to stimuli that induce long-lasting plasticity, and for memory consolidation (Costa-Mattioli et al., 2009; Richter and Klann, 2009). In AD, multiple studies have observed dysregulation of translation throughout the disease process, as indicated by changes in p70 S6 kinase 1, eukaryotic initiation factors 2 alpha (eIF2α) and 4E (eIF4E) phosphorylation (An et al., 2003; Chang et al., 2002; Ferrer, 2002; Li et al., 2004; Lourenco et al., 2013; Ma et al., 2013), as well as by decreased ribosomal function (Ding et al., 2005; Hernández-Ortega et al., 2016; Langstrom et al., 1989; Sajdel-Sulkowska and Marotta, 1984). Curiously, altered expression of ribosomal RNAs and ribosomal protein-coding mRNAs in the hippocampus appear even prior to the onset of symptoms and the development of hallmark AD pathologies (Ding et al., 2005; Hernández-Ortega et al., 2016). Furthermore, ribosomal dysfunction is specific to cortical areas that show the greatest atrophy in later stages of the disease, including the inferior parietal lobule and superior middle temporal gyri (Ding et al., 2005).

Although alterations in the level of *de novo* protein synthesis in AD have been implied for several years, the mechanistic details involved in this dysregulation remain largely unknown (Buffington et al., 2014). When examined in APP/PS1 mutant mice, which express familial AD-associated mutations in the amyloid precursor protein (APP_swe_) and presenilin-1 genes (PS1_ΔE9_) (Ma et al., 2013), an approximately 30% decrease in *de novo* protein synthesis was observed in aged symptomatic APP/PS1 mice compared to wild-type littermates (Ma et al., 2013). Elucidating the identity of these dysregulated proteins has become an important focus of research, and two independent studies of transgenic mice that experimentally model different aspects of AD pathology have used *in vivo* metabolic labelling to isolate *de novo* synthesized proteins for subsequent mass spectrometry analysis (Evans et al., 2019; Ma et al., 2020). The use of a less complex system, such as live isolated hippocampal slices, offers a different resolution and a narrower timeframe for this snapshot of the *de novo* proteome.

Herein, we used BONLAC, a combinatorial approach of stable isotope labelling by amino acid tagging (SILAC) and biorthogonal noncanonical amino acid tagging (BONCAT), and mass spectrometry to compare *de novo* protein synthesis in acute hippocampal slices from asymptomatic and symptomatic APP/PS1 mutant mice for the identification and measurement of relative abundance of *de novo* synthesized proteins (Bowling et al., 2019; Bowling et al., 2016; Dieterich et al., 2006; Eichelbaum et al., 2012; Evans et al., 2019; Ong et al., 2002; Zhang et al., 2014). Here, we show for the first time a significant dysregulation of proteostasis in APP/PS1 mutant mice even before symptom onset, with significant alterations in networks of proteins involved in both protein degradation and synthesis. Further, in aged APP/PS1 mice that display memory impairments (Volianskis et al., 2010), we observed dysregulation of both lysosomal and mitochondrial proteins, as well as components of the ribosome. Together, these findings suggest that alterations in the synthesis of proteins involved in a variety of critical cellular pathways are impaired early in the AD process and likely underlie the deterioration of neuronal function, leading to loss of memory.

## Results

### Aging-related downregulation of hippocampal protein synthesis in APP/PS1 mutant mice

Previous studies using puromycin and azidohomoalanine (AHA) incorporation to label newly synthesized proteins in mice modeling aspects of AD pathology demonstrated a reduction in *de novo* protein synthesis (Beckelman et al., 2019; Evans et al., 2019; Ma et al., 2013; Ma et al., 2020), and the translation efficiency of polyribosomes isolated from the mild cognitive impairment (MCI) and end-stage AD brain is reduced by >60% in affected cortical areas (Ding et al., 2005). We first confirmed that histopathological changes are observed in the brain of asymptomatic vs symptomatic APP/PS1 mice by performing immunohistochemistry against A*β* in brain sections including the dorsal hippocampus. As expected, we found a robust increase in the number and size of amyloid deposits in the brains of symptomatic APP/PS1 mice when compared to asymptomatic APP/PS1 mice (Figure 1A-C). Using BONCAT labeling, we observed a ∼20% decrease in the level of *de novo* protein synthesis in symptomatic (12+ months old) APP/PS1 mice compared to asymptomatic APP/PS1 mice (3-5 months-old; Figure 1D). Collectively, these findings support and extend previous work conducted in our laboratory, which highlighted a significant decrease in *de novo* protein synthesis between symptomatic wild type and APP/PS1 mice using SUnSET, another non-radioactive method for tagging and monitoring new protein synthesis (Ma et al., 2013). Therefore, using BONLAC labeling we proceeded to identify changes in the *de novo* proteome in the APP/PS1 mutant mice before and after symptom onset (Figure 1E).

**Figure 1.**
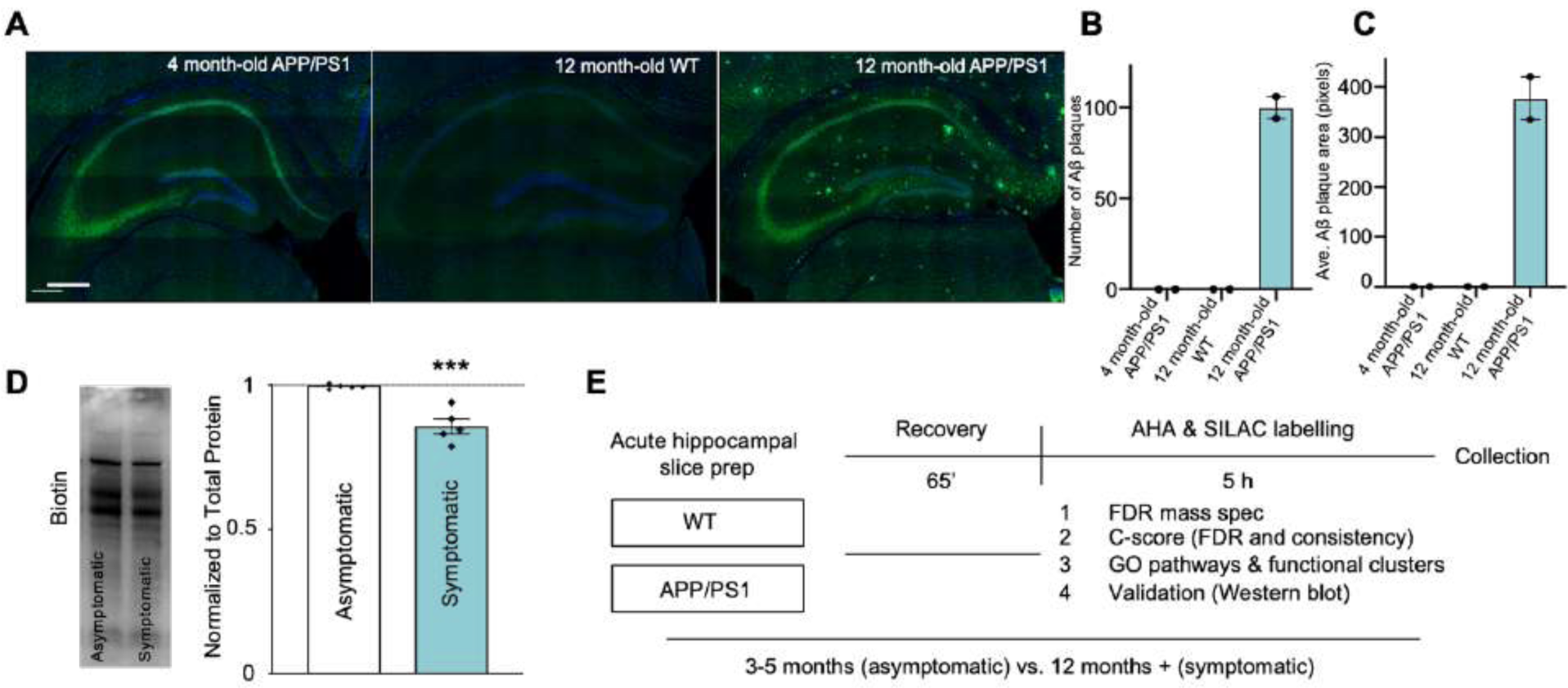
BONLAC-mediated labelling of *de novo* protein synthesis in the hippocampus of young and aged wild-type and APP/PS1 mice. (**A)** Representative immunofluorescent images showing the progressive deposition of amyloid beta plaques in the hippocampus of symptomatic 12 month-old APP/PS1 mice, but not asymptomatic 4 month-old APP/PS1 or 12 month-old wild-type (WT) mice (Blue = DAPI, Green = Aβ). **(B)** Quantification of the number of Aβ plaques in the hippocampus of asymptomatic APP/PS1 mice, 12 month-old WT and symptomatic 12 month-old APP/PS1 mice. **(C)** Quantification of the average area of the Aβ plaques observed in these groups (n = 1-2 mice per group for **(B)** and **(C)**). **(D)** *De novo* synthesized protein in 3-5 month-old asymptomatic vs. 12+ month-old symptomatic APP/PS1 mouse hippocampal slices as detected via AHA labelling, followed by biotin-alkyne click reaction and western blot (normalized to total protein (as determined via MemCode staining), expressed relative to average asymptomatic biotin signal; *n* = 5 mice/condition; statistical significance calculated using unpaired two tailed t tests: *p* = 0.0005). **(E)** Schematic showing BONLAC workflow for acute hippocampal slices from WT and APP/PS1 mice. Following labelling, slices from one APP/PS1 and one age-matched WT mouse are pooled for subsequent processing.

### Large-scale proteomic dysfunction is observed early in pathology development in the hippocampus of APP/PS1 mice

In order to identify candidate proteins of interest, we used BONLAC and a previously published multi-layered analysis based in coincidence detection (Bowling et al., 2019; Bowling et al., 2016). In the hippocampi of asymptomatic mice (3-5 months-old), 2510 *de novo* synthesized proteins were measured in at least one sample. 1826 proteins were measured across the majority of samples (Supplementary Figure 1), and of these 1826, 180 were differentially regulated in the APP/PS1 mice (Figure 2A; Supplementary Figure 2). The majority of differentially regulated proteins were downregulated (fold change ≤ 0.8; 107 or 5.9% of total proteins), whereas 73 proteins were upregulated (fold change ≥ 1.2; 4.0% of total proteins; Figure 2B) in the APP/PS1 mice. When the biological relevance of these changes were interrogated using DAVID and StringDB, the GO and KEGG pathways that were highlighted included the cellular process ‘Alzheimer’s disease’, as well as proteostasis-related components, gathered under the terms ‘proteasome’ and ‘ribosome’ (Figure 2C-D). These findings indicate that even in young APP/PS1 mice that do not show display memory deficits, processes previously linked to AD and critical to cell functioning are disturbed.

**Figure 2.**
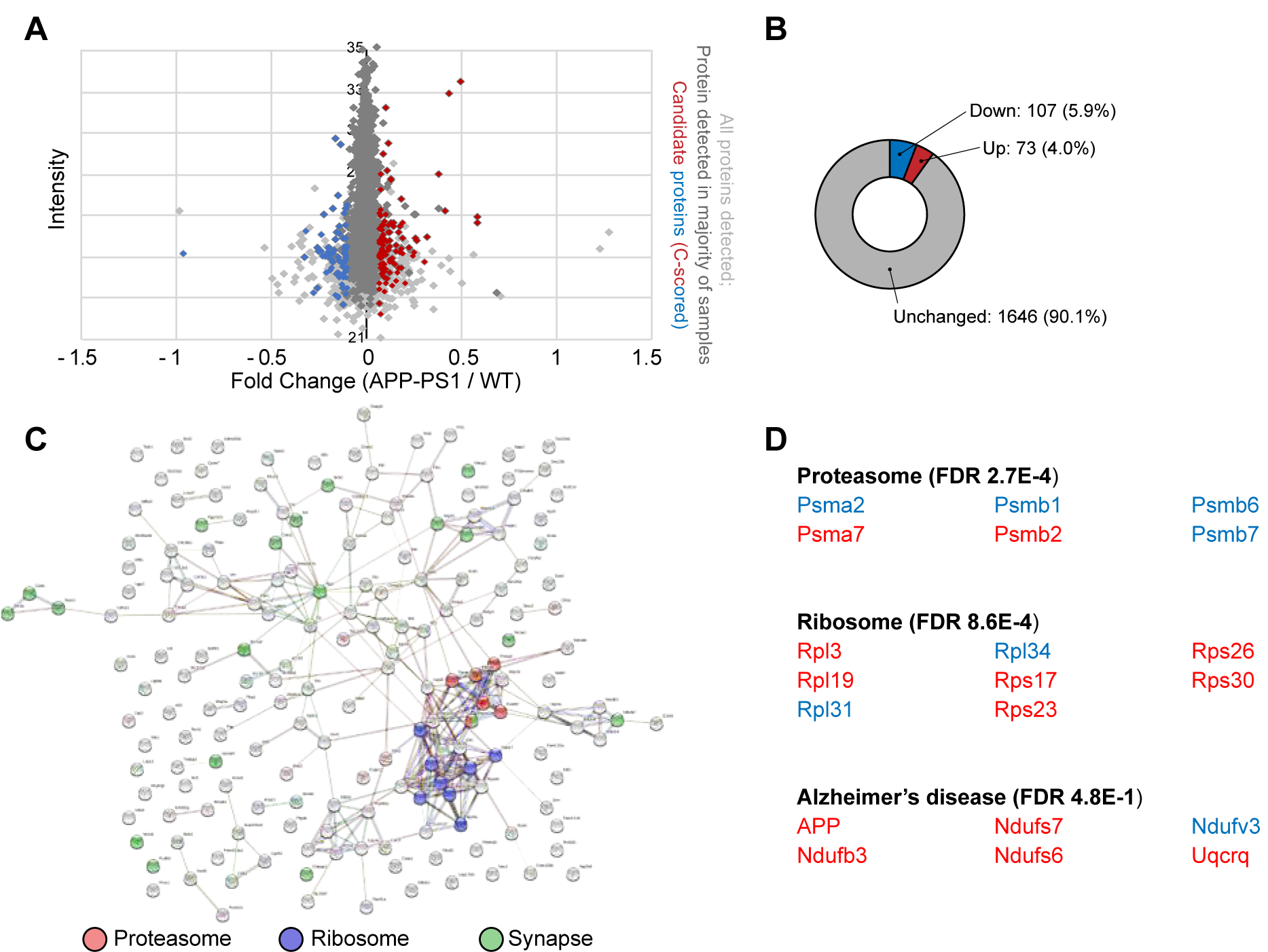
Steady state proteome differs in wild-type and asymptomatic APP/PS1 mouse hippocampus. **(A)** Fold change vs. intensity distribution plot of all newly made proteins detected in wild-type (WT) vs. 3-5 month-old APP/PS1 hippocampus (light grey), with proteins that were detected in majority of samples (>3 out of 5 samples; grey) and dysregulated candidate proteins identified by C-score screen (upregulated ≥ 20% in red, downregulated ≤ 20% in blue). **(B)** Doughnut plot showing the proportion of detected proteins that were downregulated (fold change < 0.8, blue), upregulated (fold change > 1.2, red) or unchanged in the majority of asymptomatic APP/PS1 mice compared to WT littermates. **(C)** String diagram depicting enriched gene ontology groups. **(D)** Top functional clusters in 3-5 month-old APP/PS1 mice compared to WT using DAVID. Red = upregulated proteins. Blue = downregulated proteins.

### Alterations in the *de novo* proteome correspond with pathologies in symptomatic APP/PS1 mice

Following these investigations in asymptomatic APP/PS1 mice, we investigated the impact of aging and the development of AD-like pathology on the *de novo* proteome. As expected from the decreased rate of translation apparent in AHA incorporation levels (Figure 1D), fewer newly synthesized proteins were observed in APP/PS1 mice aged ≥ 12 months compared to younger mice. In the hippocampi of symptomatic mice, 2065 proteins were measured at least once (Supplementary Figure 3), 855 proteins were measured across the majority of samples, and 113 proteins were consistently altered in the APP/PS1 mice (Figure 3A; Supplementary Figure 4). Proteins whose synthesis was reduced by 20% compared to age-matched wild-type mice made up 2.7% of the consistently detected proteins (23 proteins), while BONLAC detected 90 proteins with increased *de novo* synthesis in the symptomatic APP/PS1 mice (10.6%; Figure 3B). Through GO analysis via DAVID and visualization using StringDB, we observed several functional clusters (Figure 3C), with the top hits including ‘Alzheimer’s disease’, ‘ribosome’, and ‘lysosome’ (Figure 3D). Together, these findings indicate that the dysregulation of core cellular processes relies at least partially on impaired protein synthesis, and that changes in *de novo* proteome happen even before full pathology onset.

**Figure 3.**
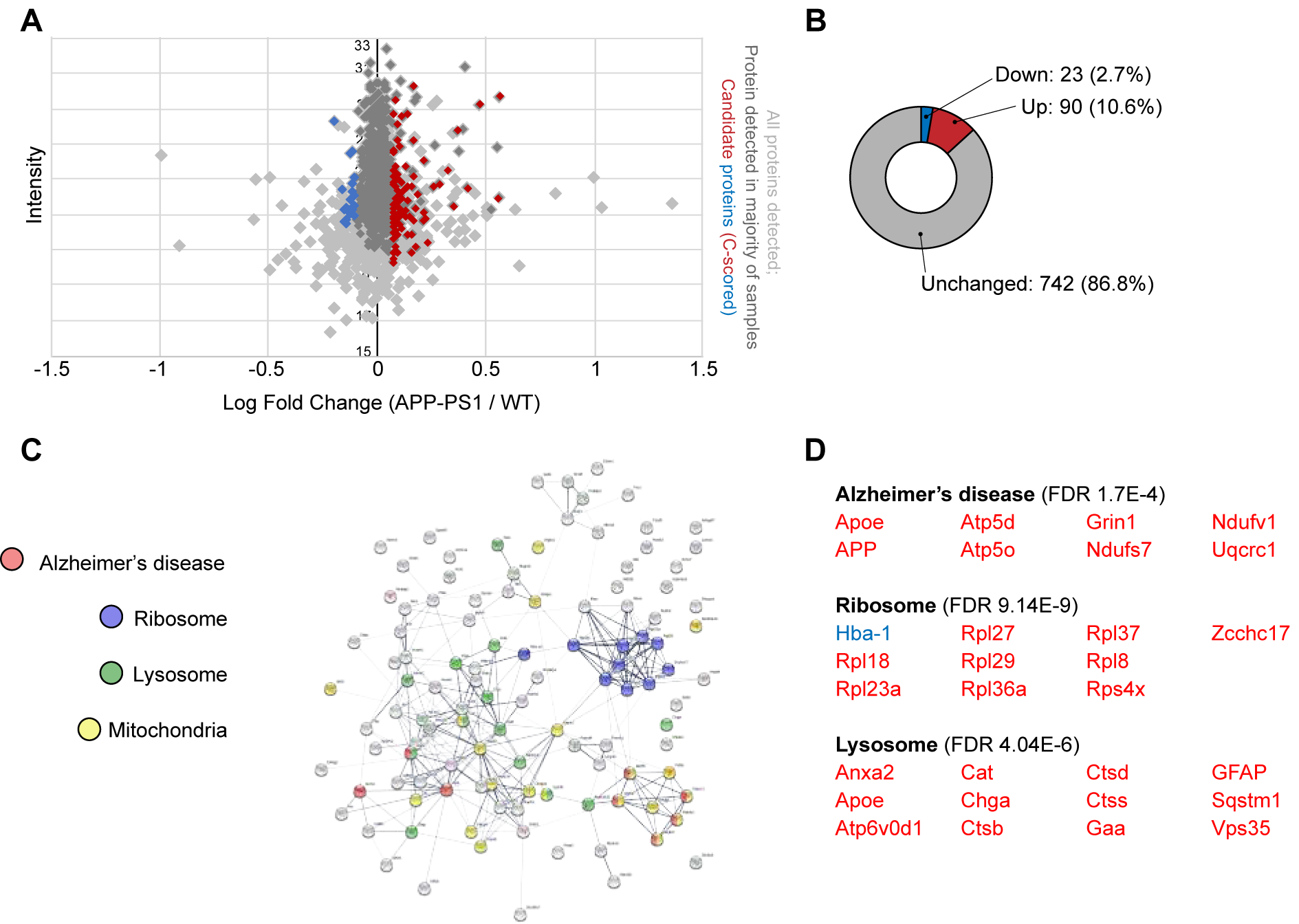
Analysis of altered *de novo* proteome highlights impaired proteostasis in symptomatic APP/PS1 mice. **(A)** Fold change vs. intensity distribution plot showing all proteins detected in 12+ month-old APP/PS1 mouse hippocampal slices vs. wild-type (WT) littermates using BONLAC (light grey). Dark grey overlay depicts proteins that were consistently detected in the majority of samples (>4/7), and candidate proteins identified by C-score screen as upregulated (fold change ≥ 1.2) shown in red, while downregulated proteins (≤ 0.8) are labelled blue. **(B)** Doughnut plot indicating the majority of proteins detected in symptomatic mice are not altered in APP/PS1 mice (742; 86.8% of proteins detected). 23 proteins are downregulated compared to WT mice (2.7% of proteins show a fold change >0.8, blue) and 90 proteins are upregulated in APP/PS1 mice (10.6% of proteins show a fold change <1.2, red). **(C)** String diagram showing enriched gene ontology networks. **(D)** DAVID-identified functional clusters in 12+ month-old APP/PS1 mice. Red = upregulated proteins. Blue = downregulated proteins.

### Dysregulation of the synthesis of BONLAC-identified candidates is reflected in protein expression

Next, to understand whether these changes in *de novo* protein synthesis corresponded to altered protein expression, we validated candidates from several enriched clusters using western blotting, comparing changes in expression between APP/PS1 and wild-type littermates. In order to validate differences from the same group used for proteomic studies, we reserved BONLAC-labeled acute hippocampal slices from animals prior to sample preparation for mass spectrometry analysis, and instead lysed the tissue for western blot analysis (Figure 4A). Alongside APP, which was selected as a positive control (Radde et al., 2006), other AD-associated proteins known to be involved in lysosome-mediated protein degradation (cathepsin-d; ctsd (Cataldo et al., 1995; Hamazaki, 1996)) and synaptic plasticity (neuromodulin; GAP-43 (Bogdanovic et al., 2000; de la Monte et al., 1995)) were probed. Samples were also probed for the expression of the large 60S ribosomal proteins Rpl13 and Rpl18. As shown in Figure 4B and Figure 4C, the *de novo* synthesis and the total expression of APP were increased in both age groups of APP/PS1 mice, as expected. Neither the synthesis nor the expression of Ctsd protein were altered in asymptomatic APP/PS1 mice compared to wild-type littermates. However in aged, symptomatic mice, a similar trend was found in both BONLAC and total protein levels. In contrast to younger mice, the older APP/PS1 mice also showed significantly reduced synthesis and total expression of the synaptic protein GAP-43, which corresponds with a known decline in synaptic density at this age (Alonso-Nanclares et al., 2013).

**Figure 4.**
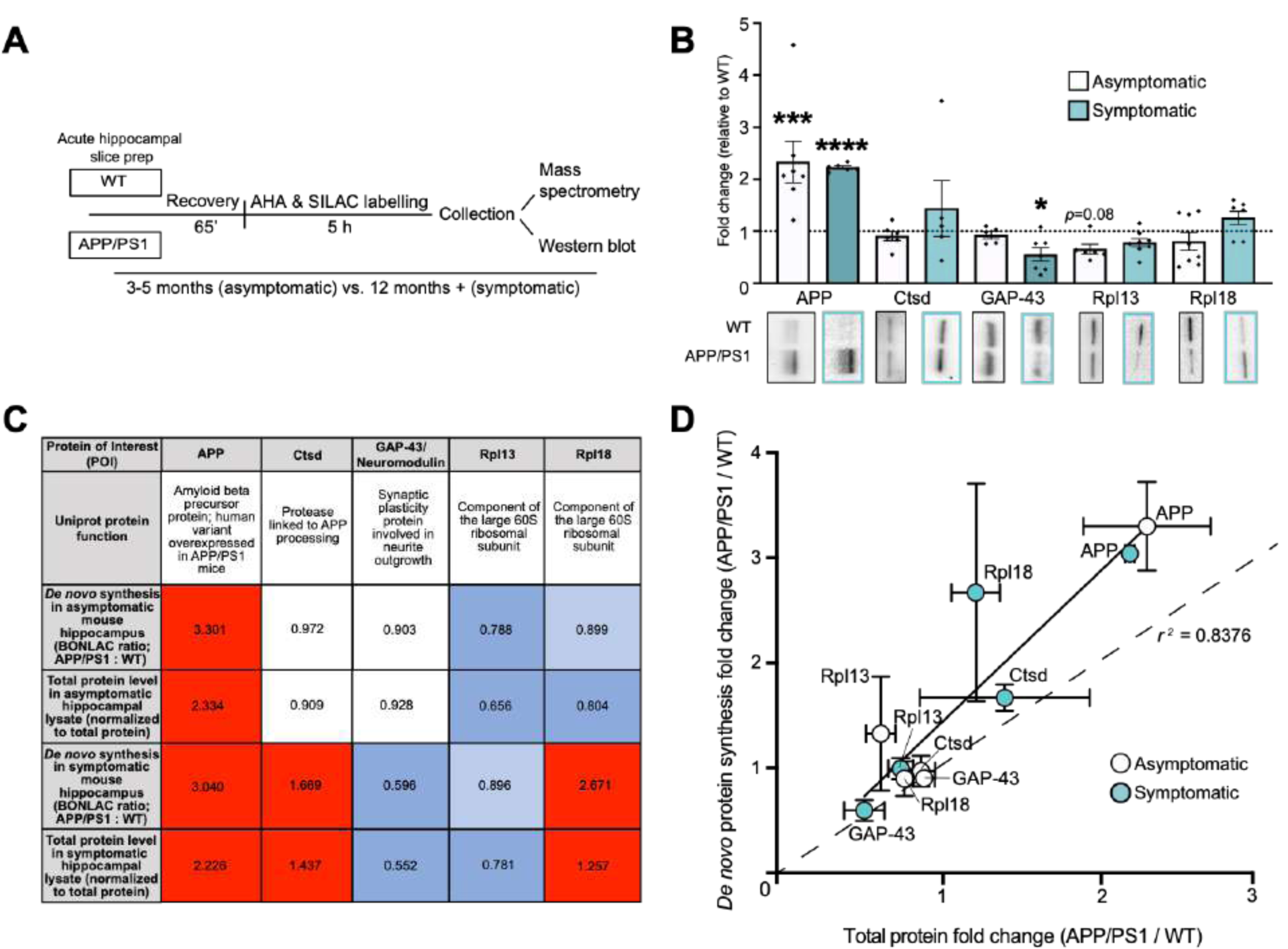
Candidate proteins identified with BONLAC screen of *de novo* proteome exhibit altered expression levels in APP/PS1 mice. **(A)** Schematic of workflow for slice allocation for mass spectrometry and western blot protein validation. **(B)** Western blot quantitation of candidate proteins selected from BONLAC screen in asymptomatic (white bars) and symptomatic (teal bars) mice, normalized to total protein (as assessed by MemCode) and expressed relative to wild-type (WT; *n* = 5-8 mice); representative western blots showing selected protein levels in WT and APP/PS1 mouse hippocampal lysates (asymptomatic: black box; symptomatic: teal box). **(C)** Comparison table showing average fold change in *de novo* synthesis of candidate proteins in asymptomatic and symptomatic APP/PS1 vs WT littermates as identified by BONLAC (asymptomatic n=5; symptomatic n=7) compared to change in total expression quantified by western blot. Vibrant color indicates value reached cutoff (+/- 20%; >0.8 or < 1.2); pale color indicates near threshold (within 5%); white indicates no change. Statistical significance calculated using unpaired two tailed t tests; * = *p* ≤ 0.05, *** *p* ≤ 0.001, **** *p* ≤ 0.0001. **(D)** Linear regression showing correlation between average *de novo* synthesized protein fold-change (as determined by BONLAC) vs. average total protein fold-change (as determined via western blot) of candidate proteins in the APP/PS1 vs. WT hippocampus before (white circles) and after (teal circles) pathology development (*r* ^*2*^ value = 0.8376, figure shows mean +/- SEM (vertical error bars: BONLAC; horizontal error bars: western blot); *p* = 0.0002).

The large 60S ribosomal subunit protein Rpl13 also was examined after BONLAC labelling indicated synthesis of this protein was decreased in APP/PS1 mutant mice both before and after the appearance of symptoms. Western blot analysis confirmed that there was a non-significant trend in the reduction in expression of this ribosomal protein in both groups of APP/PS1 mice. The expression of an additional ribosomal protein, Rpl18, was investigated as BONLAC labelling indicated the synthesis of this protein varied throughout the development of pathology in the APP/PS1 mice. In asymptomatic mice, both newly synthesized and total expression of Rpl18 were decreased at trend level relative to wild-type mice, whereas both measures were increased in aged symptomatic APP/PS1 mice. Next, we performed a linear regression analysis between the BONLAC and western blot fold changes (APP/PS1 vs. wild type) to evaluate whether a linearity existed between the values. Linear regression analysis highlighted a significant correlation between detection of the candidate protein in the *de novo* analysis vs. candidate protein expression in the total protein fraction (Figure 4D). Together, these results indicate that altered synthesis of proteins detected with BONLAC are correlated with changes in global protein expression.

### Cluster analysis of commonly synthesized proteins in both asymptomatic and symptomatic APP/PS1 mice highlights key AD pathways

Following the corroboration of BONLAC candidates by western blotting, we further investigated the overall patterns of protein synthesis regulation observed in the APP/PS1 mice. Although examination of significant changes (as detected by C-score analysis) reveal statistically relevant alteration of the *de novo* proteome, it is noteworthy that smaller fluctuations in expression (fold change < 20%: labelling ratio > 0.8 or < 1.2) can also provide information as to whether a specific cell process is altered. Therefore, we compared all 791 newly synthesized proteins identified in both asymptomatic and symptomatic APP/PS1 mice and their wild-type littermates. To investigate whether any biological pathways were differentially affected, we conducted hierarchical clustering analysis using Cytoscape. Several key clusters were observed (Figure 5, Supplementary Table 1), closely associated with the KEGG pathways ‘Alzheimer’s disease’ (Figure 5 Cluster 5; Supplementary Figure 9) and ‘protein processing in the endoplasmic reticulum’ (Figure 5 Cluster 1; Supplementary Figure 5). The GO terms ‘synapse’ (Figure 5 Cluster 2; Supplementary Figure 6), and ‘synaptic vesicle cycle (Figure 5 Cluster 3; Supplementary Figure 7) were enriched in this analysis, as were the GO terms ‘axo-dendritic transport’ (Figure 5 Cluster 4; Supplementary Figure 8), ‘myelin sheath’ (Figure 5 Cluster 5; Supplementary Figure 9), ‘mitochondrial part’ and ‘electron transfer activity’ (Figure 5 Cluster 6; Supplementary Figure 10).

**Figure 5.**
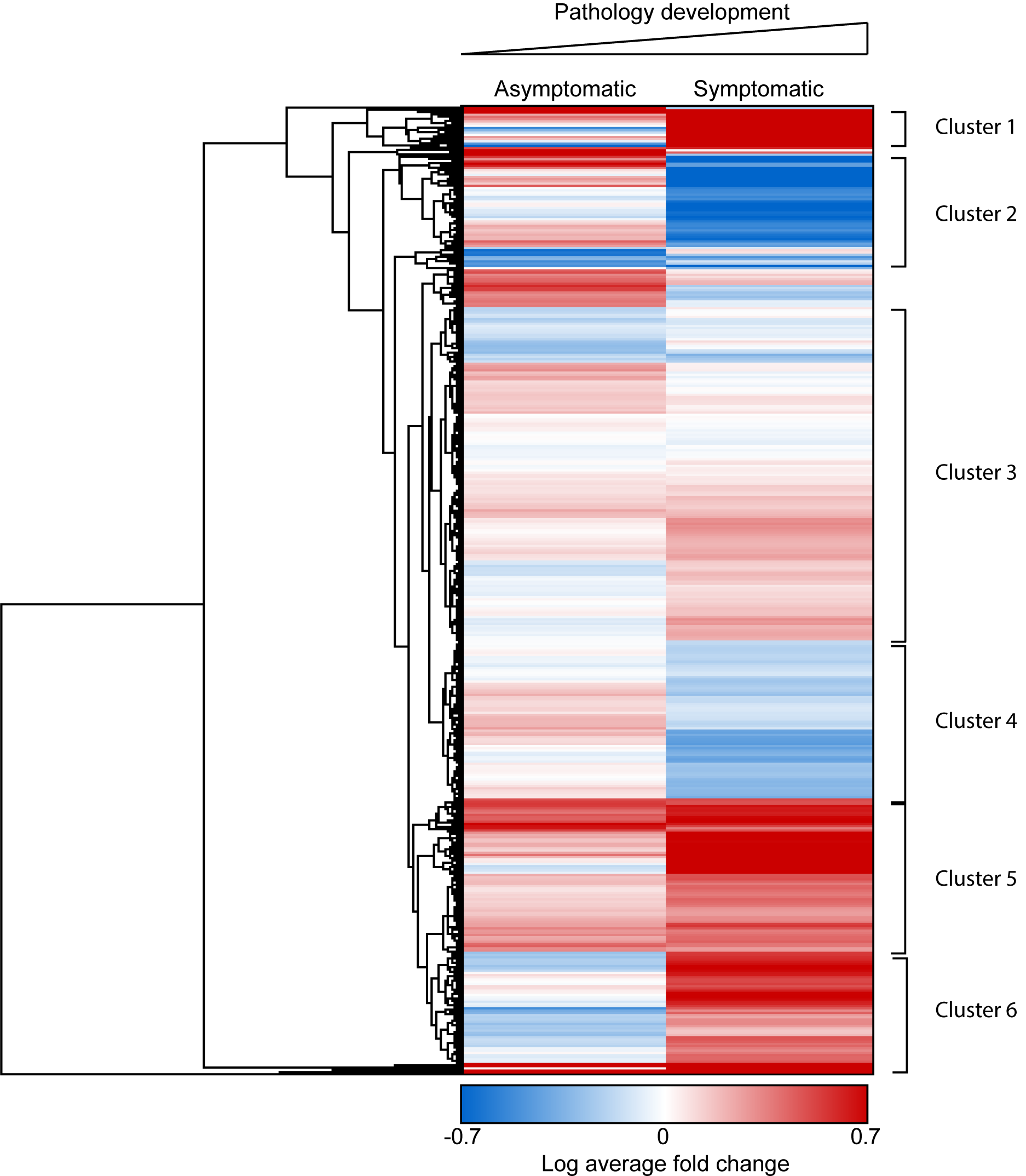
The biological pathways affected by dysregulated hippocampal protein synthesis vary throughout pathological progression in APP/PS1 mice. Hierarchical clustering heatmap reveals similarities and divergence in the mean-normalized log protein fold change of the 791 proteins detected in both asymptomatic (∼4 months-old) and symptomatic (12+ months-old) APP/PS1 mice relative to wild-type (WT) littermates. GO analysis evidenced 6 major clusters being significantly modified, which are: 1) protein processing in the endoplasmic reticulum (FDR 3.2E-04); 2) synapse (FDR 5.6E-07); 3) synaptic vesicle cycle (FDR 2.77E-06), membrane trafficking (FDR 6.24E-11) and synapse (FDR 4.82E-29); 4) axo-dendritic transport (FDR 4.86E-08) and glycolysis/gluconeogenesis (2.56E-05); 5) myelin sheath (5.17E-18) and Alzheimer’s disease (8.16E-07); and 6) mitochondrial part (FDR 2.54E-08) and electron transfer activity (FDR 4.02E-05). FDR generated by StringDB algorithms. Red = upregulated proteins. Blue = downregulated proteins. White = unchanged protein expression.

### Dysregulation of ribosomal protein synthesis is a common feature in APP/PS1 mice throughout the development of symptoms

As described above, C-score detection of candidates generated a list of proteins that were highly dysregulated in the APP/PS1 hippocampus (+/- 20% compared to wild-type). When these dysregulated candidate proteins were compared between age groups, only 31 proteins were significantly dysregulated in both the asymptomatic and symptomatic APP/PS1 mice relative to age-matched wild-type littermates (Figure 6A). The majority of these proteins showed either an increased level of synthesis at both ages (11/31) or reversed expression (decreased synthesis in asymptomatic APP/PS1 mice and increased synthesis in symptomatic mice, or vice versa; 15/31) in comparison to wild-type mice. Only five proteins showed consistently decreased synthesis in the APP/PS1 mice, regardless of age (Figure 6B). Identification of the proteins consistently dysregulated in the APP/PS1 mice highlighted a significant cluster of ribosomal proteins, three of which are components of the small 40S ribosomal subunit, and six of which are components of the large 60S ribosomal subunit (Figure 4C). Taken as a whole, these results support the notion that alterations in the expression of ribosomal proteins is an early feature of AD pathology, and may well play a role in protein synthesis dysfunction through the course of disease.

**Figure 6.**
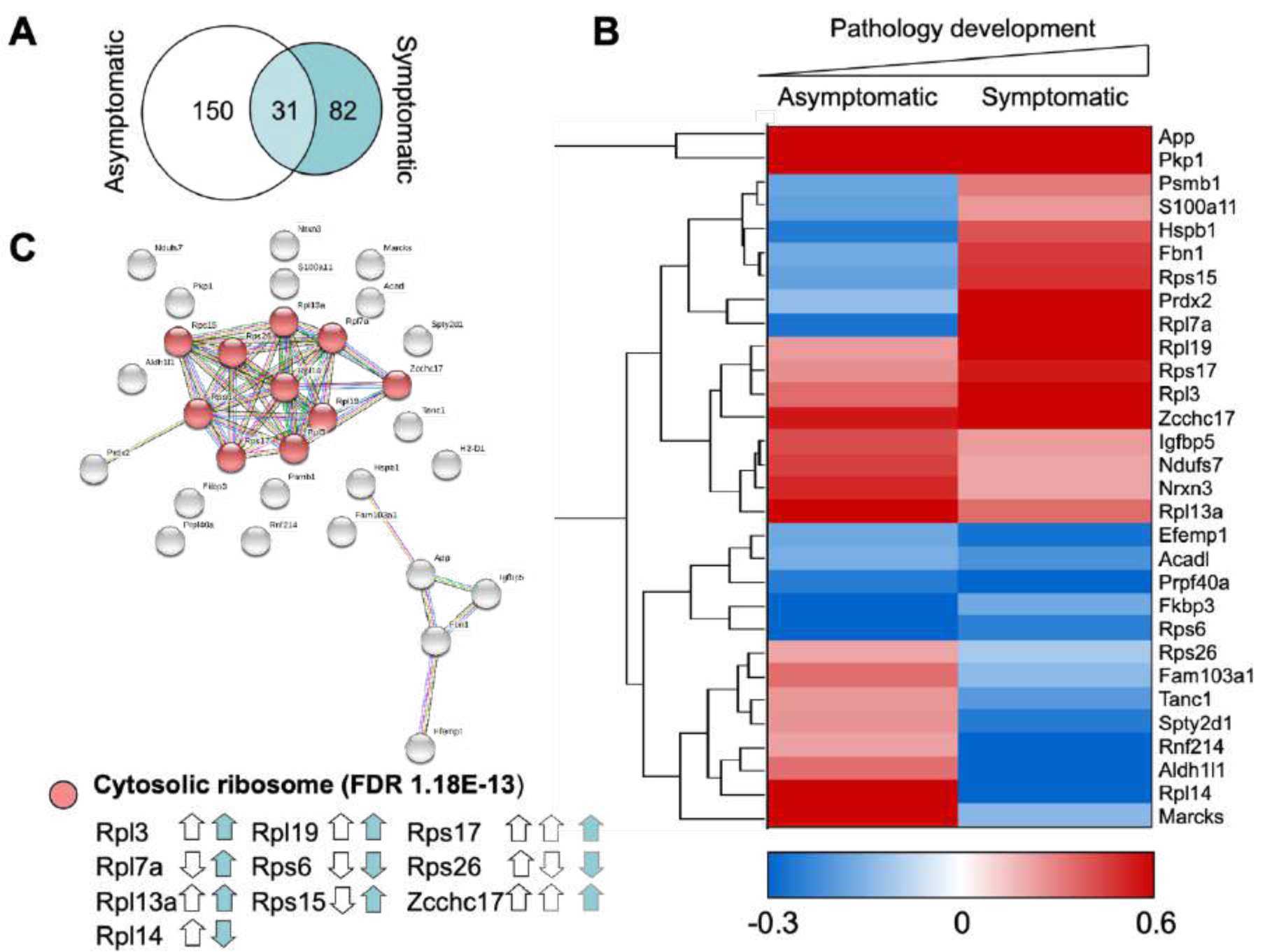
Steady state proteome is predominantly distinct throughout the development of symptoms. **(A)** Minimal overlap is observed between proteins that are dysregulated (as detected by C-score rank algorithm) in the hippocampi of 3-5 month-old asymptomatic vs. 12+ month-old APP/PS1 mice. **(B)** Hierarchical clustering of log fold changes reveals the majority of proteins which are dysregulated in both asymptomatic and symptomatic APP/PS1 mice compared to WT littermates do not show the same trend. Red = upregulated proteins. Blue = downregulated proteins. White = unchanged protein expression. **(C)** String diagram revealing an enrichment of proteins in the GO network ‘Cytosolic ribosome’ in APP/PS1 throughout the disease process. White arrows = direction of regulation in asymptomatic mice. Teal arrows = direction of regulation in symptomatic mice.

## Discussion

Dysregulated proteostasis underlies many of the hallmark pathologies of AD. Both the deposition of Aβ in the form of plaques and accumulation of hyperphosphorylated tau-containing tangles suggest impaired protein degradation systems, and indeed, dysfunctional proteolysis has been observed in AD brains (Ciechanover and Kwon, 2015). Further, translation is impaired throughout the AD process [10-17] and *de novo* protein synthesis is decreased in mouse lines that model various aspects of AD pathology [14, 23, 62]. The identity of the proteins that are differentially synthesized before and after AD symptom onset in the human brain are unknown. Interestingly however, ribosomal function was shown to be impaired in the brains of individuals diagnosed with mild cognitive impairment (Ding et al., 2005).

Recently, the *de novo* proteome of mouse lines that model aspects of AD have become a focus of study. Decreased *de novo* protein synthesis was observed in neurons with high levels of hyperphosphorylated tau in the K369I (K3) and rTg4510 transgenic mouse models of tauopathy and neurodegeneration (Evans et al., 2019). Moreover, and consistent with our findings, investigations using *in vivo* metabolic labelling has highlighted significant dysregulation of the synthesis of proteins involved in vesicle transportation and mitochondrial functioning prior to symptom onset in APP/PS1 mice (Ma et al., 2020). This study also investigated changes in the proteome both pre- and post-symptom onset, but the young, asymptomatic mice selected for the study (2 months-old) do not yet show hallmark Aβ deposition in the hippocampus, which begins at 3-4 months of age (Radde et al., 2006). Moreover, although impairments in hippocampal LTP and cognitive ability are apparent by 9 months of age (Gengler et al., 2010; Radde et al., 2006; Serneels et al., 2009), the age selected for the symptomatic group in this work, several studies have indicated that older mice (>12 months old) possess greater deficits in spatial learning and memory (Gruart et al., 2008; Park et al., 2006; Trinchese et al., 2004; Xu et al., 2013).

In the present study, we used BONLAC labelling in acute hippocampal slices from ∼4 month-old asymptomatic and ∼12 month-old symptomatic APP/PS1 mice and their wild-type littermates to isolate and identify alterations in the *de novo* hippocampal proteome. The use of animals at these ages both supports the recent *in vivo* work described above (Ma et al., 2020), and extends the coverage of the *de novo* proteome investigations throughout pathology development in this model. Decreased synthesis was observed in aged APP/PS1 mice, as represented by lower AHA incorporation and fewer proteins identified in the mass spectrometry screen, as anticipated from prior work indicating an age-dependent decrease in translation in this AD model (Ma et al., 2013). At both timepoints, the vast majority (>85%) of measured proteins were synthesized consistently regardless of genotype, further illustrating the tight regulation of the proteome required for cell functioning. However, defined patterns of dysregulation were observed in APP/PS1 mice relative to unaffected littermates.

In young APP/PS1 mice that do not yet show symptoms, we observed a specific pattern of dysregulated protein synthesis, resulting in predominately decreased translation of proteins involved in a series of pathways critical to cell functioning. Of note, proteins involved in both protein degradation (via GO term ‘proteasome’) and protein synthesis (via GO term ‘ribosome’) showed dysregulated synthesis in the asymptomatic APP/PS1 hippocampus relative to wild-type littermates. In fact, the dysfunctional production of components of the proteasome could have a negative effect on ribosomal function and general protein synthesis, as an interplay between impaired proteasomal function and translation has previously been described (Ding et al., 2005). Dysregulated synthesis of synaptic proteins also were highlighted at this time point, which could underlie the decreased synaptic density previously observed in 4 month-old APP/PS1 mice (Smith et al., 2009), as well as the impaired synaptic transmission observed in APP/PS1 mice at 5 months of age (Zhang et al., 2005). Thus, our findings may represent a critical window in which modulation of translation limits synaptic changes that result in memory deficits later in disease progression.

When *de novo* protein synthesis was examined in 12+ month-old APP/PS1 mice that display pronounced AD-like pathology and memory impairment, a large number of mitochondrial proteins previously linked to AD were dysregulated. Mitochondrial structure and function is known to be compromised in human patients (Baloyannis, 2006; Hirai et al., 2001), but whether these deficits contribute to the disease or whether they are a biproduct of disease progression continues to be debated (Swerdlow, 2018). Alongside mitochondria, lysosomal protein synthesis was dysregulated in the symptomatic APP/PS1 mice. Increasing evidence suggests that lysosomal biogenesis is initially upregulated in early stages of AD (Cataldo et al., 1995), before progressive dysfunction throughout the disease process results in the accumulation of enlarged lysosomes, which fail to effectively degrade their contents (Nixon, 2017).

A final significant pathway that was observed in the analysis of dysregulated proteins following symptom onset was constituents of the cytosolic ribosome, including eight ribosomal proteins and nucleolar protein of 40 kDa (Zcchc17). This zinc ion binding protein has recently been identified as a modulator of synaptic gene expression in AD (Tomljanovic et al., 2018). Moreover, Zcchc17 is thought to be involved in ribosome biogenesis and maturation (Chang et al., 2003). Two protein components of the ribosome that were differentially regulated in the mature APP/PS1 tissue, Rpl13 (decreased BONLAC labelling compared to wild-type) and Rpl18 (upregulated in APP/PS1 mice) were selected for validation analyses in hippocampal lysates. The total expression of both proteins reflected the level of synthesis as detected in the mass spectrometry screen, indicating that these are biologically relevant changes that require further investigation. Rpl13 has recently been described as a ‘core’ ribosomal protein included in all actively translating ribosomes (Shi et al., 2017), and yet has also been shown to be dysregulated at the gene expression level in hippocampal tissue from severe AD patients (Kong et al., 2009). Although no study has yet linked alterations in Rpl18 expression with AD pathology, recent analysis has indicated dysregulation of *Rpl18* gene expression occurs early in the AD process (Martínez-Ballesteros et al., 2017).

In order to determine the full impact of APP/PS1 mutations on protein synthesis in these mice, we examined the biological pathways that exhibited changes across pathology development as detected by hierarchical cluster analysis of all proteins detected in both age groups. This list was not limited to proteins identified by C-score analysis as significantly dysregulated, and included proteins detected in the majority of samples which showed a fold change of <20%. Several functional clusters emerged, indicating pathways specific to stage of pathology development. Two clusters were specifically upregulated in the symptomatic APP/PS1 mice, those associated with the GO terms ‘protein processing in the endoplasmic reticulum (ER)’ and ‘mitochondrial part: electron transfer activity’. Increased synthesis of proteins, including Stub1/CHIP, Cat and Hsph1, which have previously been implicated in the AD process (Habib et al., 2010; Poirier et al., 2019; Singh and Pati, 2015), indicate that the unfolded protein response (UPR), the physiological response to the ER stress, might be chronically activated later in disease. This finding is consistent with our previous studies showing that reducing the expression of PERK, an eIF2*α* kinase that triggers downregulation of protein synthesis in response to UPR activation, could restore hippocampal plasticity and memory deficits in symptomatic APP/PS1 mice (Ma et al., 2013).

Mitochondrial dysfunction is known to be an early event in both APP/PS1 mice (Chen et al., 2019) and human AD brain pathology (Hauptmann et al., 2009; Moreira et al., 2007; Nunomura et al., 2001), and is believed to play a role in the synaptic loss that occurs early in the disease process (Du et al., 2012). Downregulated synthesis of mitochondrial proteins, specifically the oxidative phosphorylation-associated proteins Ndufs5, Ndufa12, Cox4i1 and Cox6b in the asymptomatic APP/PS1 mice could underlie the decreased electron transport chain capacity observed in this and other models of AD-like pathology (Bo et al., 2014; David et al., 2005).

Functional clusters that exhibited downregulation in the aged APP/PS1 mice compared to the asymptomatic cohort were also observed. The first cluster was associated with the GO term ‘synapse’, which may be a function of the decreased spine density previously observed in symptomatic APP/PS1 mice (Smith et al., 2009). Proteins found in this hub included those involved in glutamatergic signaling, known to be impaired in both the human AD brain and APP/PS1 mice (Minkeviciene et al., 2008). In addition, a cluster corresponding to ‘glycolysis and gluconeogenesis’ was decreased in the aged APP/PS1 hippocampal *de novo* proteome. Decreased glucose metabolism has been observed in the human AD brain using radioactive glucose labelling and PET scanning, and correlates well with AD-associated pathologies (Landau and Frosch, 2014; Markesbery, 2010; Martins et al., 2018).

Although the cluster analysis described above examined general trends, we also conducted focused analysis using C-score-mediated detection of dramatic changes in *de novo* protein synthesis, where candidate proteins were selected for further analysis if they showed a fold-change of >20% in either direction in APP/PS1 mice vs. wild-type littermates. Comparison of proteins selected in this manner in asymptomatic and symptomatic mice revealed that although few proteins were dysregulated in both groups, one third of these proteins were identified as ribosomal subunits. The ribosome is composed of four ribosomal RNA molecules and 80 ribosomal proteins (Yoshihama et al., 2002), which together form the small and large ribosomal subunits that function together to translate mRNAs into proteins. Recent examination of *de novo* hippocampal protein synthesis following prolonged in vivo metabolic labelling revealed dysregulated synthesis of Rps3a (Ma et al., 2020). Moreover, dysregulated expression of genes encoding several of the ribosomal proteins observed in our study (Rpl7, Rps6, Rps17 and Rps26) has previously been described in the AD brain (Hernández-Ortega et al., 2016). Notably, decreased synthesis of ribosomal proteins is a common feature in both amyloid and tauopathy mouse models, as a recent *de novo* proteomic study using K3 and rTg4510 models of tauopathy revealed dysfunctional translation of these proteins (Evans et al., 2019).

The protein content of functional ribosomes was long thought to be largely homogenous, but recent studies have indicated sub-stoichiometric inclusion of ribosomal proteins in actively-translating ribosomes in mammalian cells (Shi et al., 2017; Slavov et al., 2015). In neurons, active remodeling of ribosomal protein content has been observed in response to axonal stimulation (Shigeoka et al., 2019). Notably, ribosomal protein gene expression is regulated throughout the aging process in the brain (Ximerakis et al., 2019), indicating that the dynamic regulation of ribosomal components may play a role in cellular adaption to aging. Further, deficiency in particular ribosomal proteins results in hypersensitivity to deficits in the protein degradation system (McIntosh et al., 2011). Importantly, the protein constituents of the actively translating ribosome appear to confer a level of preference for which mRNAs are translated, suggesting a further level of translational control (Shi et al., 2017). Together with the aforementioned studies, our findings make a strong case for further detailed investigations into the dysregulation of ribosomal protein synthesis and its subsequent consequences in AD.

One important consideration when using BONLAC labeling is that the primary output is the ratio of medium to heavy isotope labelling, which is used to generate the fold-changes that permit relative quantitation. A caveat to using this system in AD (in which translation is known to be impacted) is that if a protein is not synthesized at all during the labeling window, or synthesis levels are below detection in one sample within the pair, a ratio will not be generated. As we have observed, both here (Figure 1A) and previously (Ma et al., 2013; Ma et al., 2020), a significant decline in gross *de novo* protein synthesis in the APP/PS1 mouse (Ma et al., 2013; Ma et al., 2020), this inherent bias may have led to an exaggerated detection of upregulated vs. downregulated proteins in the symptomatic APP/PS1 hippocampal samples. Bioinformatic comparison of the *de novo* hippocampal proteome generated in this, and recent *in vivo* analysis (Ma et al., 2020), with the basal proteome could reveal proteins that were not synthesized over the labelling period and were thus precluded from BONLAC ratio generation. Further by combining BONLAC labelling and TMT sample multiplexing, the number of missing peptide quantification values in each sample is greatly reduced, providing deeper coverage of the *de novo* proteome (Zecha et al., 2019). The incorporation of this novel approach into subsequent studies promises to reveal new insights into the cellular processes that accompany pathology progression in the APP/PS1 mouse.

In summary, here we have illustrated a compelling picture of dysregulation of the *de novo* proteome in the APP/PS1 mouse model of AD-like amyloidy, both before and after the appearance of symptoms. Following validation of several candidate proteins to support the relevance of the changes observed in the mass spectrometry screen, we conducted robust bioinformatic analysis of the newly made proteins. We observed dysregulated synthesis of proteins involved in several pathways known to be altered throughout the course of AD, including those involved in synaptic, mitochondrial and lysosomal functions, but we also observed significant disturbance of protein components of the ribosome. In light of recent research that indicates inclusion or exclusion of specific ribosomal subunits bestows selectivity to mRNA translation, the significant up- and down-regulation of ribosomal protein synthesis identified here, even prior to the development of pathology, may underly the progressive deterioration of cellular function and memory observed in this AD model, and in individuals with AD.

## Materials and Methods

### Animals

All procedure involving animals were performed in accordance with protocols approved by the New York University Animal Welfare Committee and followed the National Institutes of Health (NIH) *Guide for the Care and Use of Laboratory Animals*. All mice were house in the New York University animal facility. Mice of both sexes were used. APP/PS1 transgenic mice (B6;C3-Tg(APPswe, PSEN1/dE9)85Dbo/Mmjax) and their wild-type littermates were bred and maintained on C57-BL6 and B6.C3 (Jackson Labs) backgrounds. Mice were housed with their littermates in groups of two to three animals per cage and kept on a 12-hour regular light/dark cycle, with food and water provided ad libitum. All genotypes were verified by polymerase chain reaction. Mice were used at an age of either 3-5 months or 12-15 months.

### Immunohistochemistry

Mice were deeply anesthetized with ketamine (150 mg/kg) and transcardially perfused with 0.1M PBS followed by 4% PFA before brains were removed and post-fixed for 48 hours. 40 μm free-floating coronal sections containing the hippocampus were cut using a Leica vibratome, collected and stored at 4°C in 0.01% sodium azide until use. Sections were permeabilized in 0.5% Triton X-100 (15 min) prior to blocking (5% normal goat serum, 0.1% Triton X-100; 1 h). Slices were incubated overnight at 4°C in rabbit anti-amyloid beta antibody (1:200; clone 6E10, Enzo Life Sciences), followed by Alexa-488-labelled goat anti-rabbit secondary antibody (1:500; RT, 1.5 h). Slices were mounted using ProLong Gold Antifade Mountant with DAPI. Tile-scan z stack images (10-15 optical sections depending on slice thickness) were collected using an SP8 confocal microscope (Leica) with a 20X magnification lens and Leica LASX software, with laser intensity, smart gain and offset maintained throughout the experiment. Images were processed using ImageJ 2.0.0 using the Bio-Formats importer plug-in. Plaque number was manually quantified using the Cell Counter plugin (*n* = 1-2 animals/group, 1 dorsal hippocampal slice/animal).

### AHA and SILAC Dose

AHA and SILAC labels were used as previously described (Bowling et al., 2016). AHA was purchased from Click Chemistry Tools, AZ, USA, and SILAC amino acids (13C6-15N2-lysine, 13C6-15N4-arginine (Lys8/Arg10) and D4-lysine/13C6-arginine (Lys4/Arg6)) were obtained from Cambridge Isotope Laboratories, MA, USA and used at previously described concentrations (Bowling et al., 2016; Zhang et al., 2014). Assignment of SILAC labels to experimental conditions (wild-type or APP/PS1) was alternated between biological replicates to ensure that results were not biased by labelling.

### Acute Hippocampal Slice Preparation and Incubation

400 μm transverse hippocampal slices were obtained as previously described (Chévere-Torres et al., 2012; Kaphzan et al., 2013; Santini et al., 2015). Slices were recovered and incubated in ACSF for 20 min at room temperature, then at 32°C for 45 min. AHA (1 mM) and SILAC amino acids were added to the ACSF, and slices were incubated for 5 hrs. Following the labelling period, slices were immediately flash frozen for mass spectrometry and stored at - 80 °C until use.

### BONLAC Sample Preparation for Mass Spectrometry

Samples were prepared as previously described (Bowling et al., 2016) using the Click-IT Protein Enrichment Kit (ThermoFisher Scientific). BONLAC was carried out with a minimum of five runs per condition, with each run examining hippocampal slices from one wild type and one age-matched APP/PS1 animal (*n* = 5-7 biological replicates made up of 1 APP/PS1 and 1 wild type; 5-7 animals of each age per genotype were used in total). Briefly, following labeling, equal weights of tissue from age-matched pairs of wild type and APP/PS1 animals were lysed together in buffer containing 8 M urea, 200 mM Tris pH 8, 4% CHAPS, 1 M NaCl and protease inhibitor cocktail (cOmplete, Mini, Roche; two tablets per 10 mL of lysis buffer). The lysate was sonicated before AHA-labeled proteins were covalently coupled to alkyne-tagged agarose beads using reagents provided in the kit. Beads were washed repeatedly with SDS (1% SDS, 100 mM Tris pH 8, 250 mM NaCl and 5 mM EDTA) and alkyne-bound proteins were reduced with DTT at 70 °C before alkylation with iodoacetamide protected from light at room temperature. Beads were then washed sequentially to remove non-specifically bound proteins with 100 column volume SDS wash buffer, 8 M urea, and finally with 20% acetonitrile. Bound proteins were digested on-resin with trypsin (Trypsin Gold, Mass spectrometry grade, Promega) at 37 °C overnight in 25 mM ammonium bicarbonate, and the resulting tryptic peptides were desalted using hand-packed StageTips (Rappsilber et al., 2007). Desalted peptides were dried to a small droplet, under vacuum, in a SpeedVac.

### Liquid Chromatography and Mass Spectrometry

LC-MS was conducted using a Thermo Scientific EASY-nLC 1000 coupled to a Q Exactive, High Field mass spectrometer (ThermoFisher Scientific) equipped with a nanoelectrospray ionization source. Peptide separation was achieved with a self-packed ReproSil-Pur C18 AQ 3μ reverse phase column (Dr.Maisch GmbH, Germany, 75 μm inner ID, ∼25cm long). Peptides were eluted via a gradient of 3 - 40% acetonitrile in 0.1% formic acid over 120 min at a flow rate of 250 nL/min at 45 °C, maintained with a column heater (Sonation GmbH, Germany). The mass spectrometer was operated in data-dependent mode with survey scans acquired at a resolution of 120,000 at m/z 400. Up to the top 15 most abundant precursors from the survey scan were selected with an isolation window of 1.6 Thomsons and fragmented by higher energy collisional dissociation with normalized collision energies of 27. The maximum ion injection times for the survey scan and the MS/MS scans were 60 ms, and the ion target value for both scan modes were set to 1,000,000.

### Protein Identification and Quantitation

Raw files obtained from mass spectrometry runs were processed using the MaxQuant computational proteomics platform (Version 1.5.5.1(Cox and Mann, 2008)) for peptide identification and quantitation. Fragmentation spectra were searched against the Uniprot mouse protein database (downloaded on 12/20/2017, containing 16,950 non-redundant protein entries, combined with 262 common contaminants), allowing for up to two missed tryptic cleavages. Cysteine carbamidomethylation was set as a fixed modification, and methionine oxidation, acetylation of protein N-terminal, D-4 lysine, 13C6-arginine 13C6-15N2-lysine and 13C6-15N4-arginine were used as variable modifications for the database search. The mass tolerances were set to 7 ppm for precursor, and 10 ppm for fragment respectively. A false discovery rate (FDR) of 1% was applied to both peptide and protein identifications.

### Data Availability

The raw mass spectrometry data generated during this study are available at MassIVE (Center for Computational Mass Spectrometry, University of California, San Diego) with the accession number (ftp://massive.ucsd.edu/MSV000085962/). Note to reviewers: data can be accessed prior to publication using the following details: User: MSV000085962; Password: a

### Computational Processing of BONLAC MS data

MaxQuant-normalized H/M ratios (heavy vs. medium isotopes) were used for quantitative analysis. Ratios were inverted for experiments where isotopic labelling was reversed. A custom-made *R* script was used to select protein candidates where the average ratio was above or below an arbitrary threshold of 20% (>0.8 or <1.2). Only proteins that were measured in the majority of samples and were consistently up- or down-regulated following manual examination of the dataset were selected. This method allows for non-biased selection of up- or down-regulated proteins, taking variation between replicates into account.

### Protein Identity and Interaction Interpretation

Gene Ontology analysis was performed using the Database for Annotation, Visualization and Integrated Discovery (DAVID version 6.8) (Sherman and Lempicki, 2009). Candidate proteins selected according to the methods outlined above were added to the software as a ‘gene list’, and the background was considered as all proteins previously measured in hippocampal brain slices (Bowling et al., 2016). To depict the function clustering of the data described by DAVID, the STRING database (version 11.0) was used. This online resource contains both known and predicted protein-protein interactions (Szklarczyk et al., 2019). Clusters of proteins of interest were highlighted for visualization, and FDR of GO term was noted.

For hierarchical clustering of datasets, Cytoscape (Version 3.71) was used. Datasets were uploaded to the program as lists of proteins, and STRING enrichment was conducted against the mouse genome. Fold change (average or individual APP/PS1:wild type samples, as appropriate) were uploaded against the relevant network. Hierarchical clusters were generated using the clusterMaker plugin, with pairwise average linkage and Euclidean distance metric.

### Western Blot Validation

All western blotting was carried out as previously described (Sharma et al., 2010). Briefly, hippocampal tissue reserved from samples prior to mass spectrometry processing was homogenized in lysis buffer protease and phosphatase inhibitors. Protein concentration was measured using BCA assay (Pierce). Aliquots of protein (20-40 μg) were separated using Bolt Bis-Tris gels (4-12%; Thermo Scientific) and transferred to a nitrocellulose membrane which was probed using appropriate primary and secondary antibody pairs. Membranes were washed and probed for total protein using MemCode Reversible Protein Stain (Thermo Scientific), before antibody signal was detected using chemiluminescence (GE Healthcare). Band density values were normalized to total protein as detected using MemCode. Mean band densities for samples from APP/PS1 mouse samples were normalized to corresponding samples from wild type animals.

### Antibodies

The following antibodies were used in this study: rabbit anti-APP monoclonal antibody (1:1000; ab32136, Abcam), mouse anti-cathepsin-d antibody (1:500; ab6313, Abcam), rabbit anti-GAP-43 polyclonal antibody (1:500; ab5220, Sigma), anti-Rpl13 antibody (1:1000; PA5-41715, Thermofisher), and anti-Rpl18 antibody (1:5000; ab241988, Abcam). Secondary antibodies were either goat anti-mouse IgG HRP or goat anti-mouse IgG HRP (Promega; 1:10,000) respectively.

## Acknowledgments

The authors acknowledge Dr. Francesco Longo for his assistance with hippocampal slice preparation and Dr. Heather Bowling for her contributions to the experimental design. This work was supported by NIH grants S10OD023659 and NS050276 to TN, and NS047384 and NS034007 to EK. We thank the NYU Mass Spectrometry Core for Neuroscience at the NYU School of Medicine for computing resources, bioinformatics support, and assistance with data analysis and interpretation.

## Supplementary Figures

**Supplementary Figure 1.**
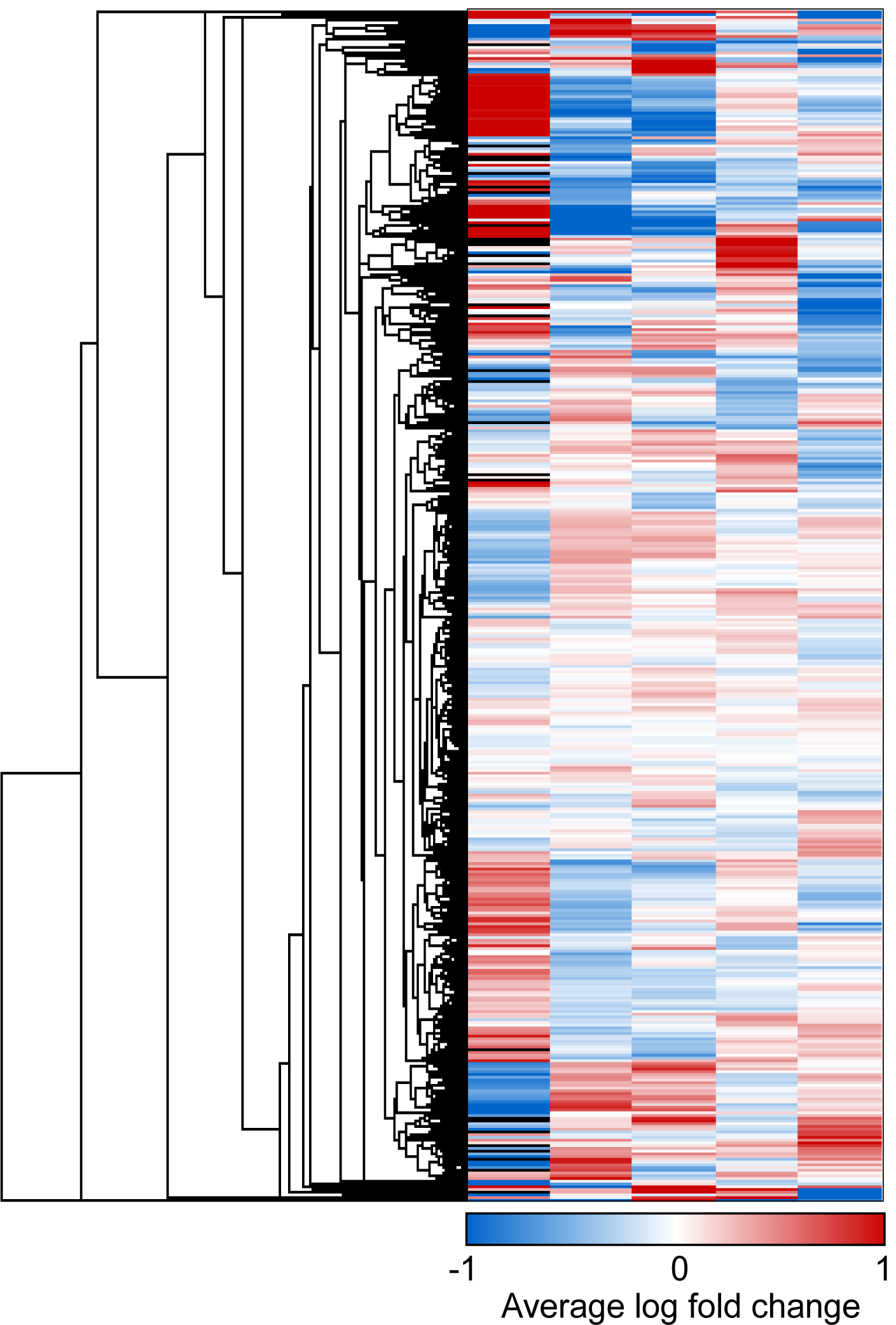
All proteins detected in the asymptomatic APP/PS1 hippocampus as detected by BONLAC. Hierarchical clustering-generated heatmap showing all protein fold-change ratios identified in at least one sample from the BONLAC screen. Red indicates proteins who show higher levels of *de novo* synthesis in ∼4 month-old APP/PS1 mice compared to wild-type (WT) littermates, while downregulated proteins are shown in blue. White = no change. Black = MaxQuant ratio not calculated.

**Supplementary Figure 2.**
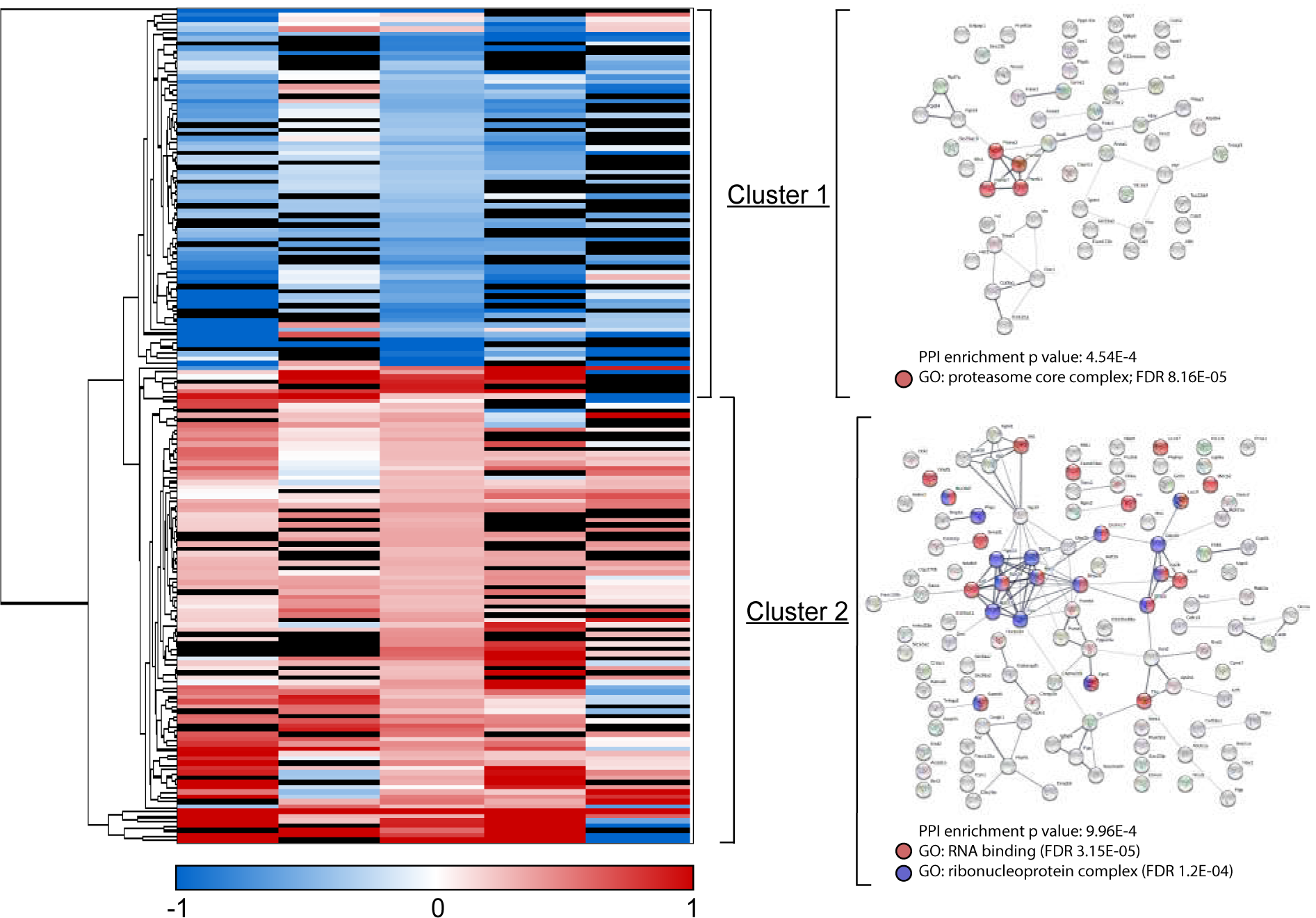
Dysregulated protein candidates in the asymptomatic APP/PS1 hippocampus as detected by BONLAC. Hierarchical clustered heatmap showing all candidate proteins identified from the BONLAC screen of ∼4 months-old APP/PS1 vs. wild-type (WT) ranked by automated C-score. Protein candidates were identified by *R* script as showing an average fold change across all samples of ± 20% (<0.8 or >1.2). Manual validation confirmed candidates were detected in the majority of samples (>3/5), and that majority of samples showed the same trend (>50% either <0.79 or >1.19). Nodes in blue indicate downregulation, while red indicates increased *de novo* synthesis in the asymptomatic APP/PS1 mice compared to WT littermates. Protein clusters are represented visually by String diagrams (right), with key pathways highlighted. White = no change. Black = MaxQuant ratio not calculated.

**Supplementary Figure 3.**
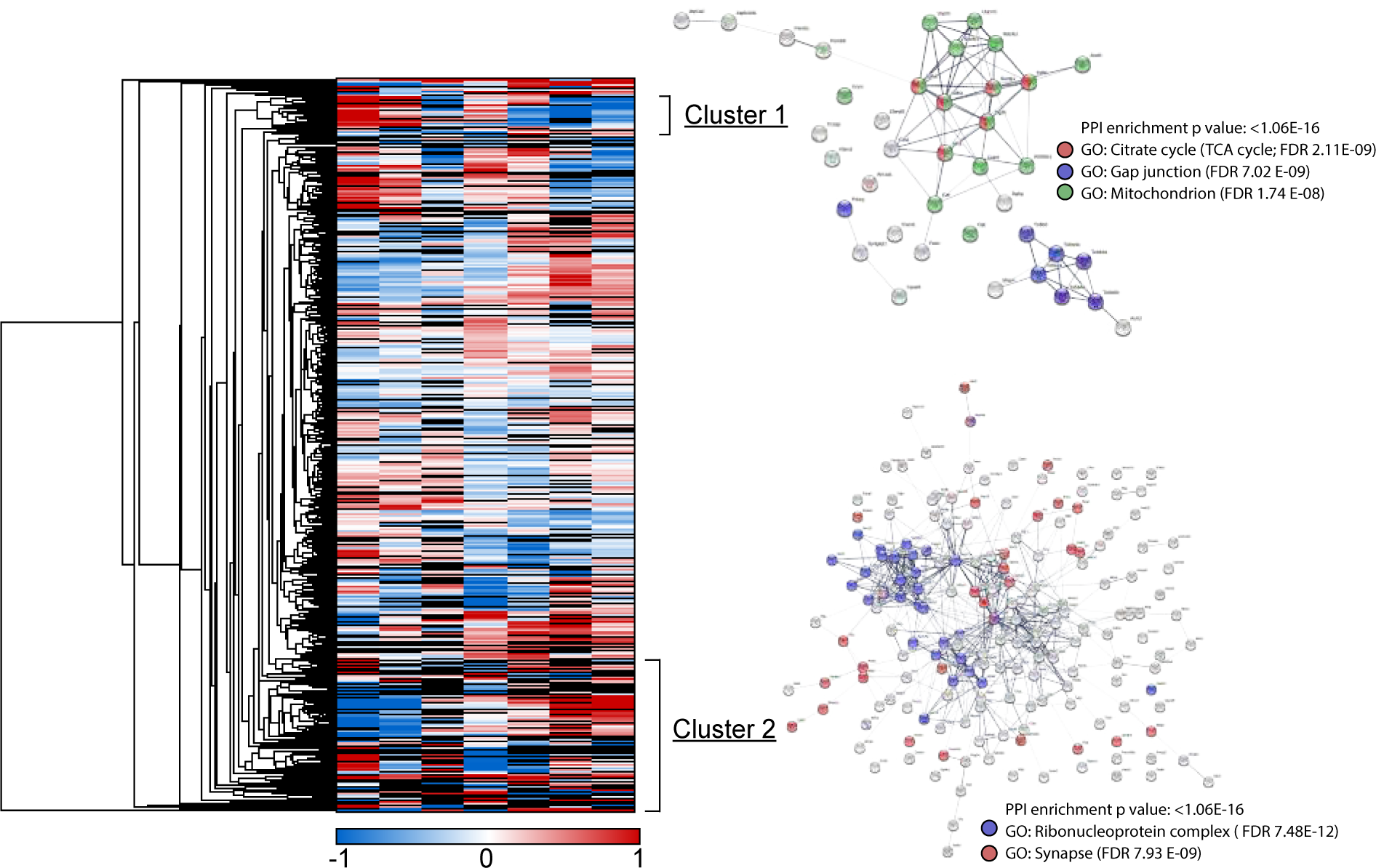
All proteins detected in the symptomatic APP/PS1 hippocampus as detected by BONLAC. Hierarchical clustering heatmap showing all protein fold-change ratios identified in the majority of samples from the BONLAC screen. Red indicates proteins which show higher levels of *de novo* synthesis in 12+ month-old APP/PS1 mice compared to WT littermates (APP is highlighted in top left), while downregulated proteins are shown in blue. String diagram of selected cluster is shown on right. White = no change. Black = MaxQuant ratio not calculated.

**Supplementary Figure 4.**
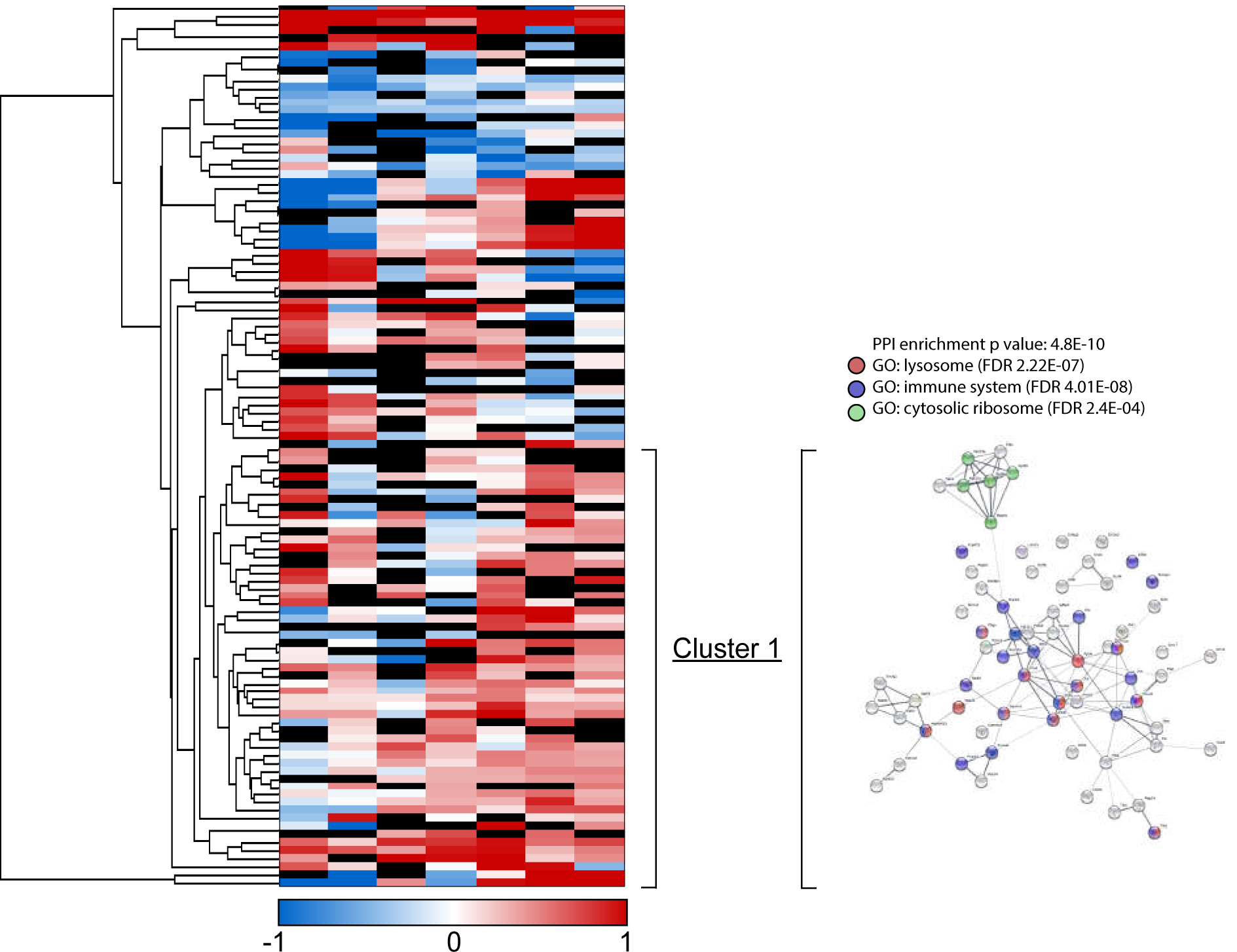
Dysregulated protein candidates in the hippocampus of symptomatic APP/PS1 mice as detected by BONLAC. Hierarchical clustered heatmap of candidate proteins identified from the BONLAC screen ranked by automated C-score. Protein candidates were identified by customized *R* script as showing an average fold change across all samples of ± 20% (<0.8 or >1.2). Manual validation confirmed candidates were detected in the majority of samples (>4/7), and that majority of samples showed the same trend (>50% either <0.79 or >1.19). String diagram representing main clusters shown on right, with key pathways highlighted. Black cells indicate absence of ratio. Red = upregulated proteins. Blue = downregulated proteins. White = no change. Black = MaxQuant ratio not calculated.

**Supplementary Figure 5.**
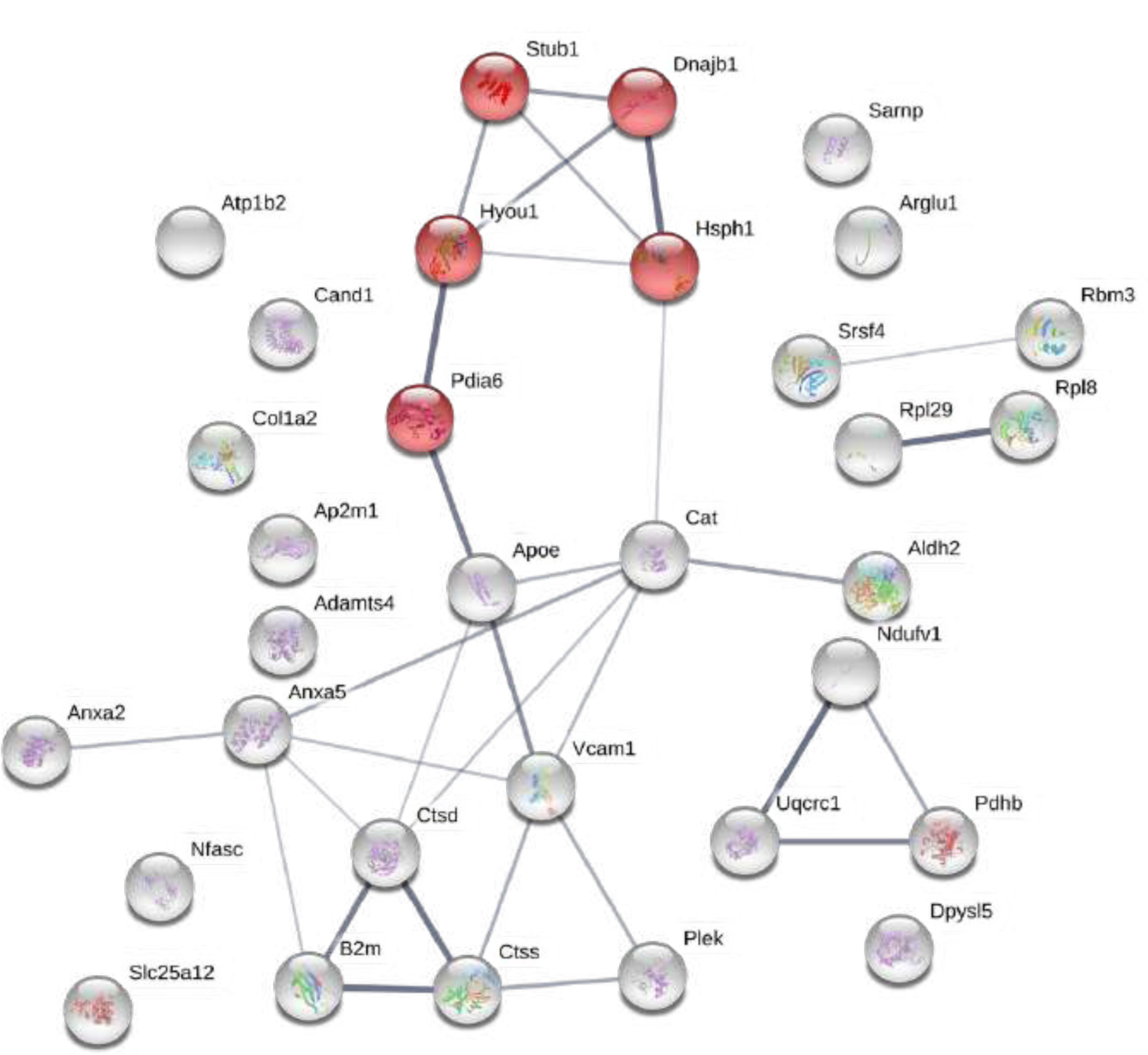
Visual depiction of proteins in Hierarchical Cluster 1 (from Figure 5). String diagram showing biological networks between the proteins identified in Cluster 1. Figure generated by StringDb; Red node = GO: Protein processing in the endoplasmic reticulum; FDR:3.2E-4.

**Supplementary Figure 6.**
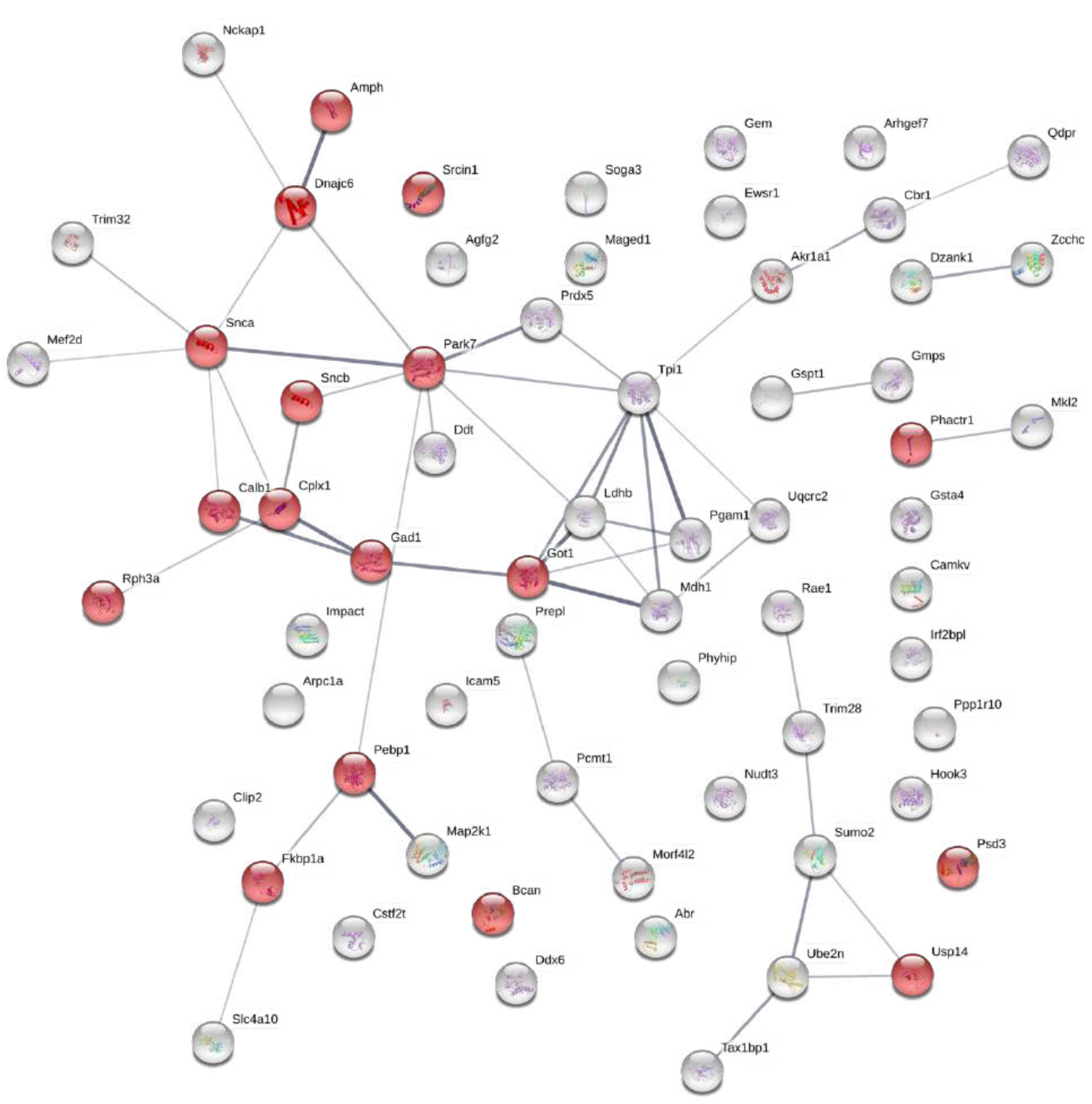
Visual depiction of proteins in Hierarchical Cluster 2 (from Figure 5). String diagram showing biological networks between the proteins identified in Cluster 2. Figure generated by StringDb; Red node = GO: Synapse; FDR:5.6E-7.

**Supplementary Figure 7.**
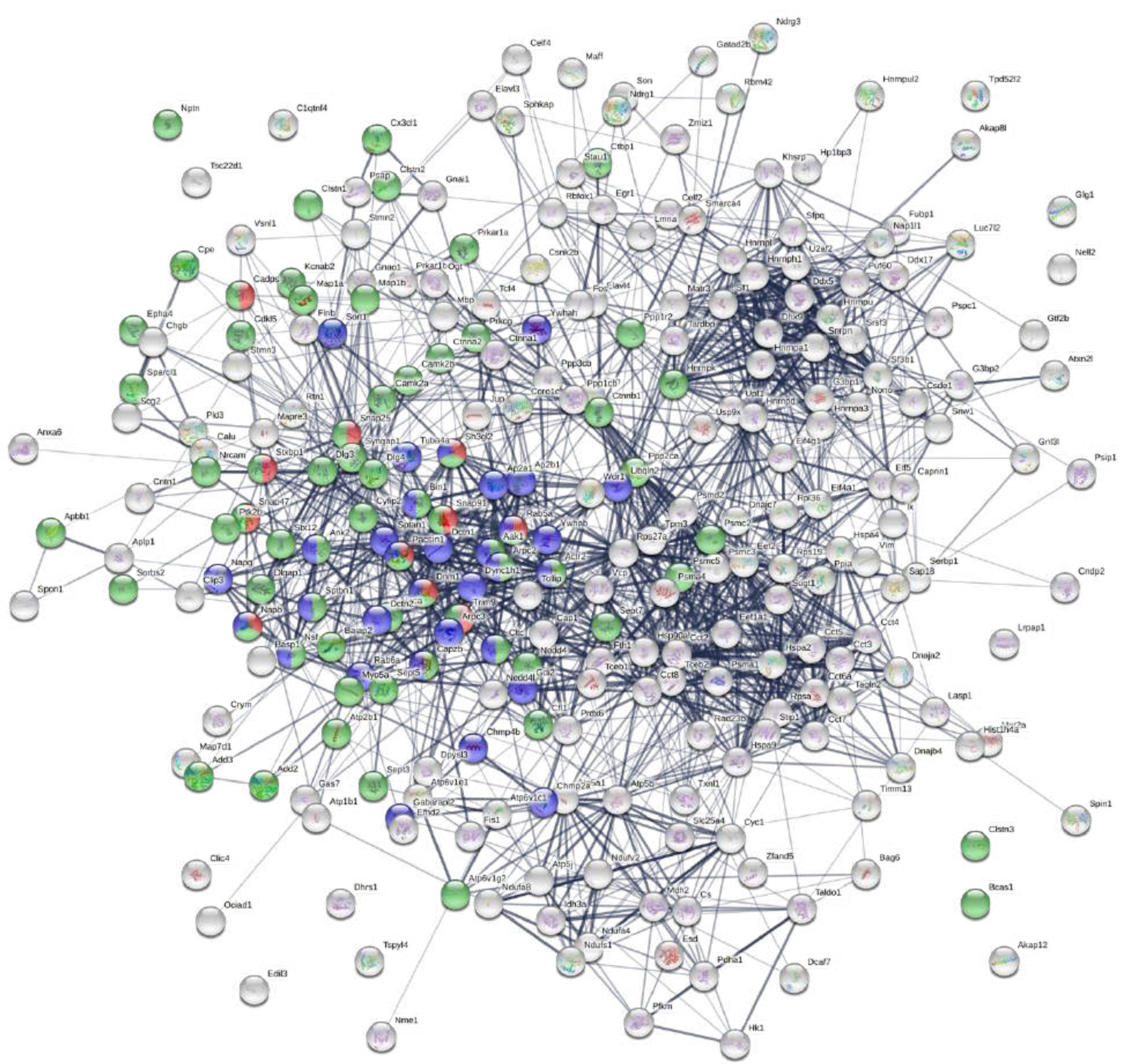
Visual depiction of proteins in Hierarchical Cluster 3 (from Figure 5). String diagram showing biological networks between the proteins identified in Cluster 3. Figure generated by StringDb; Red node = GO: synaptic vesicle cycle (FDR: 2.77E-6); Green node = GO: synapse (FDR: 4.82E-29); Blue node = GO: membrane trafficking (FDR: 6.24E-11).

**Supplementary Figure 8.**
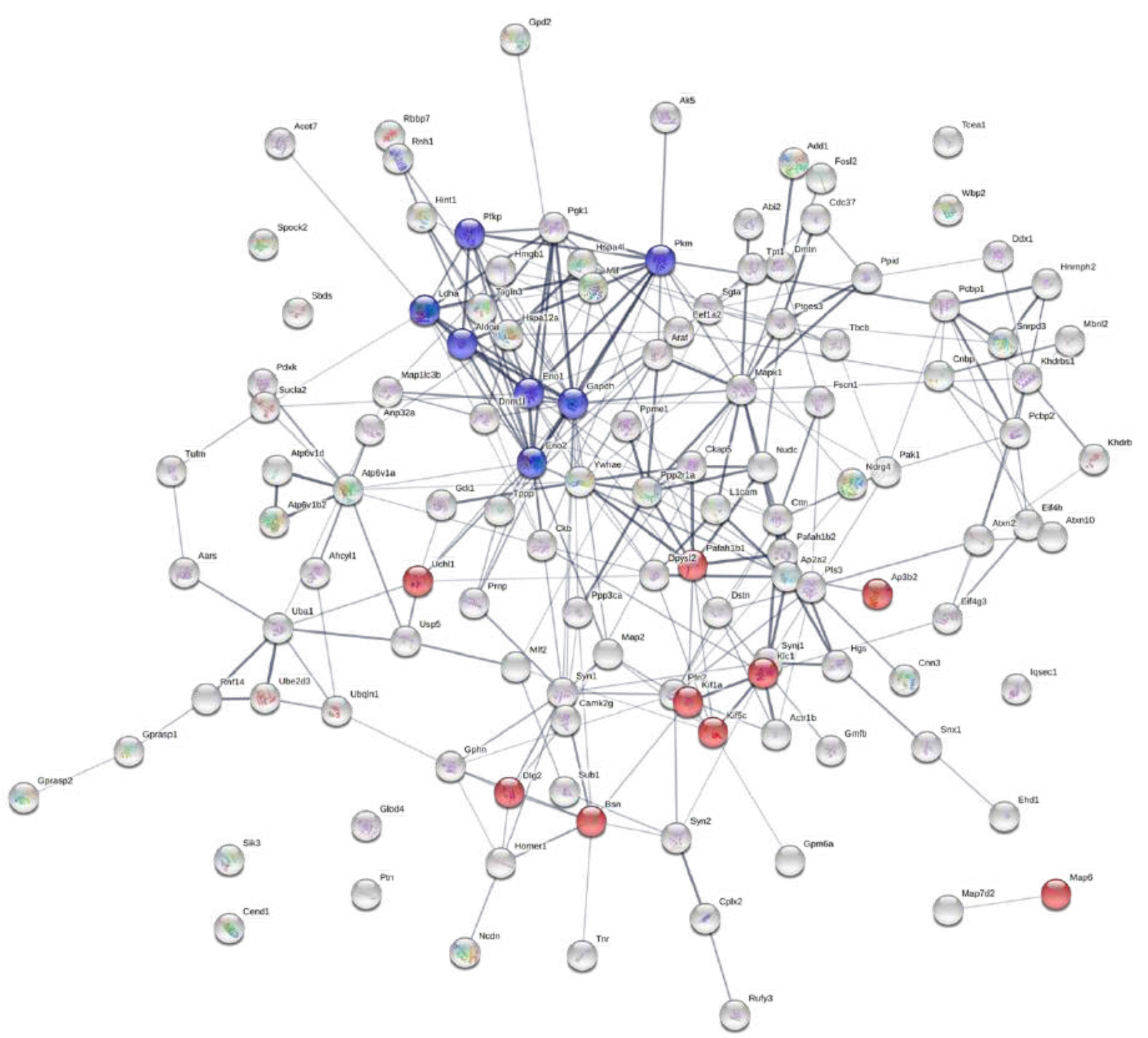
Visual depiction of proteins in Hierarchical Cluster 4 (from Figure 5). String diagram showing biological networks between the proteins identified in Cluster 4. Figure generated by StringDb; Red node = GO: Axo-dendritic transport (FDR: 4.86E-8); Blue = GO: Glycolysis/Gluconeogenesis (FDR: 2.56E-5).

**Supplementary Figure 9.**
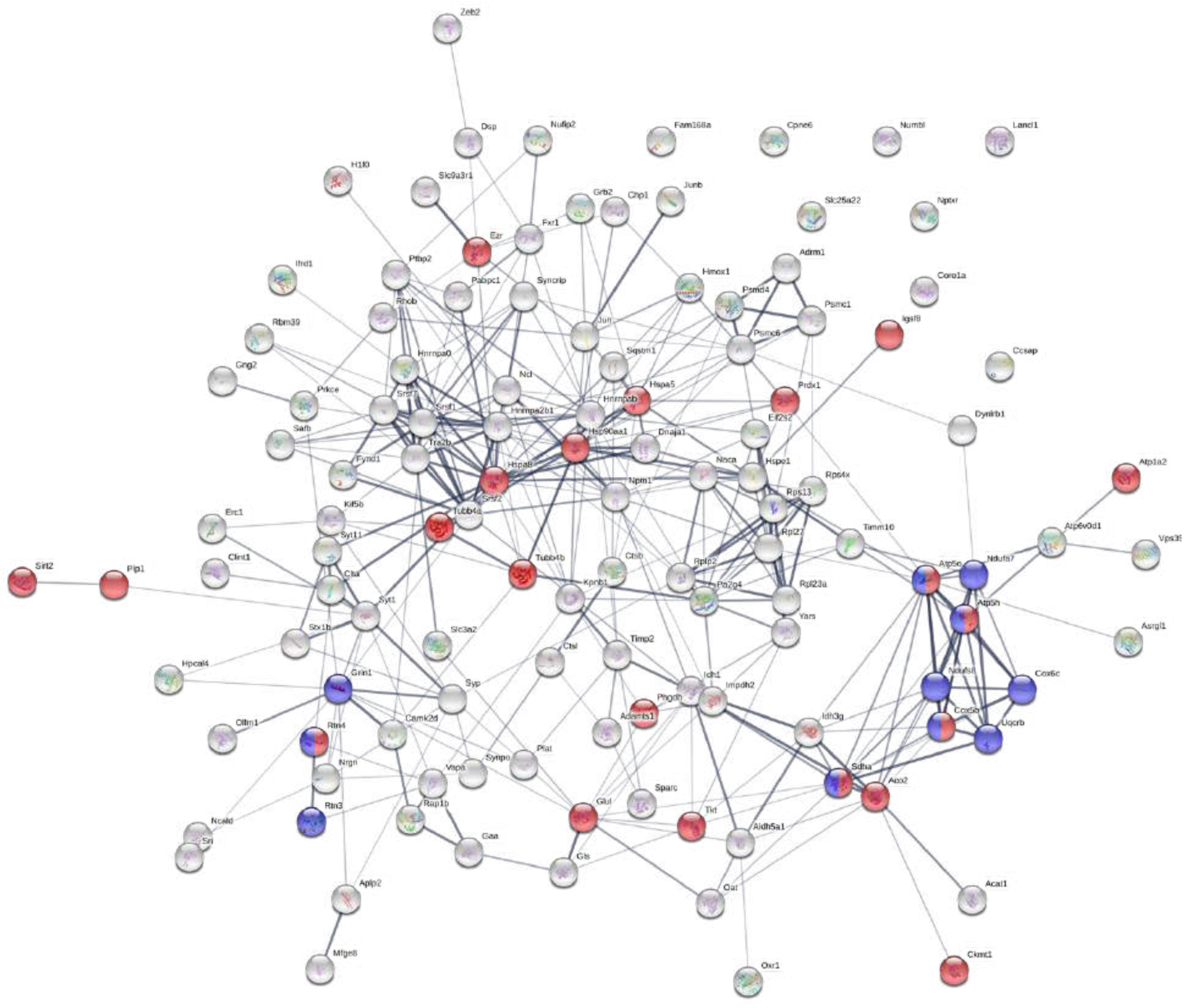
Visual depiction of proteins in Hierarchical Cluster 5 (from Figure 5). String diagram showing biological networks between the proteins identified in Cluster 5. Figure generated by StringDb; Red node = GO: myelin sheath (FDR: 5.17E-18); blue node = GO: Alzheimer’s disease (FDR: 8.16E-07).

**Supplementary Figure 10.**
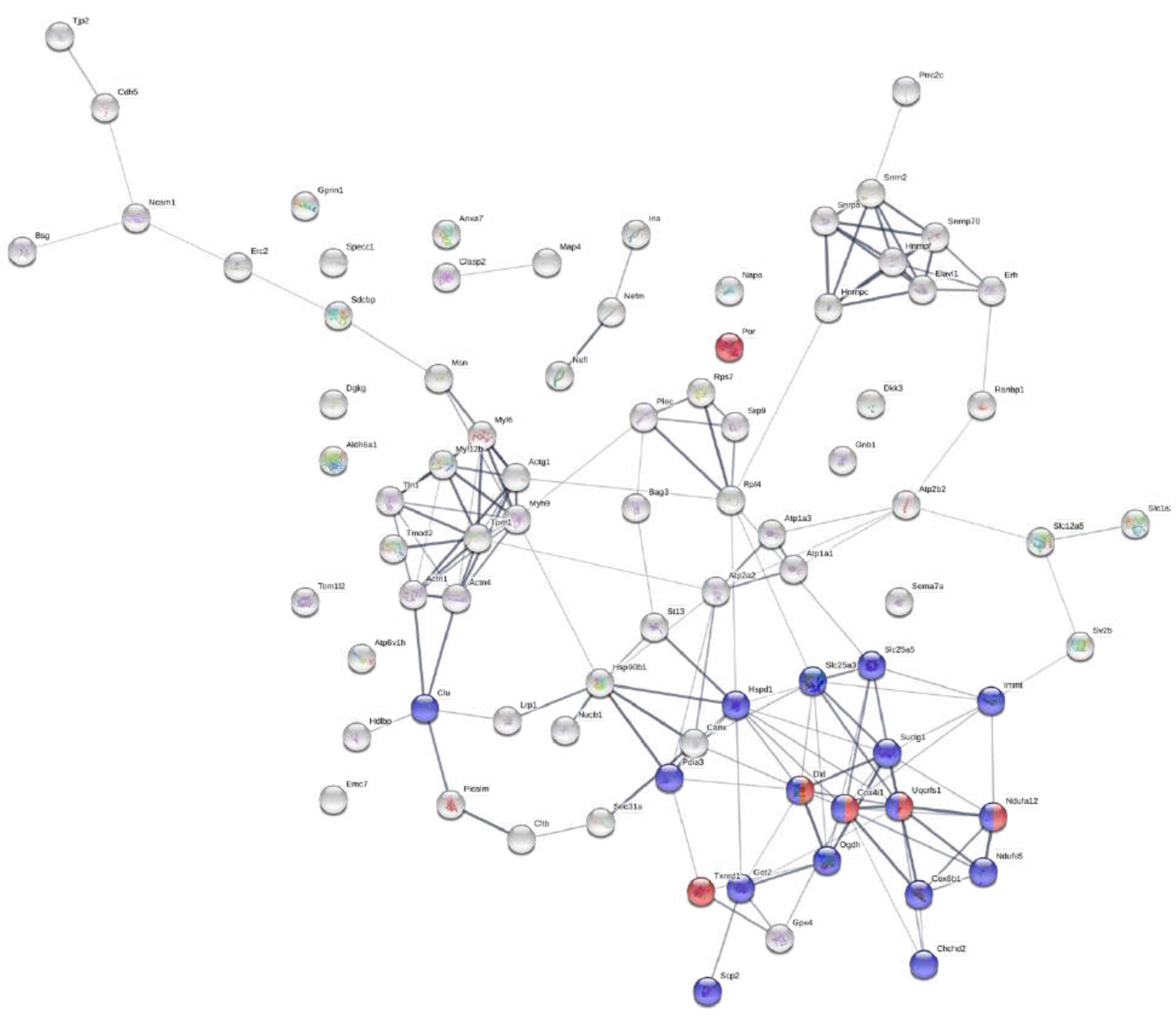
Visual depiction of proteins in Hierarchical Cluster 6 (from Figure 5). String diagram showing biological networks between the proteins identified in Cluster 6. Figure generated by StringDb; Blue node = GO: mitochondrial part (FDR: 4.22E-07); Red node = GO: electron transfer activity (FDR: 2.88E-05).

## Supplementary Table

**Supplementary Table 1.**
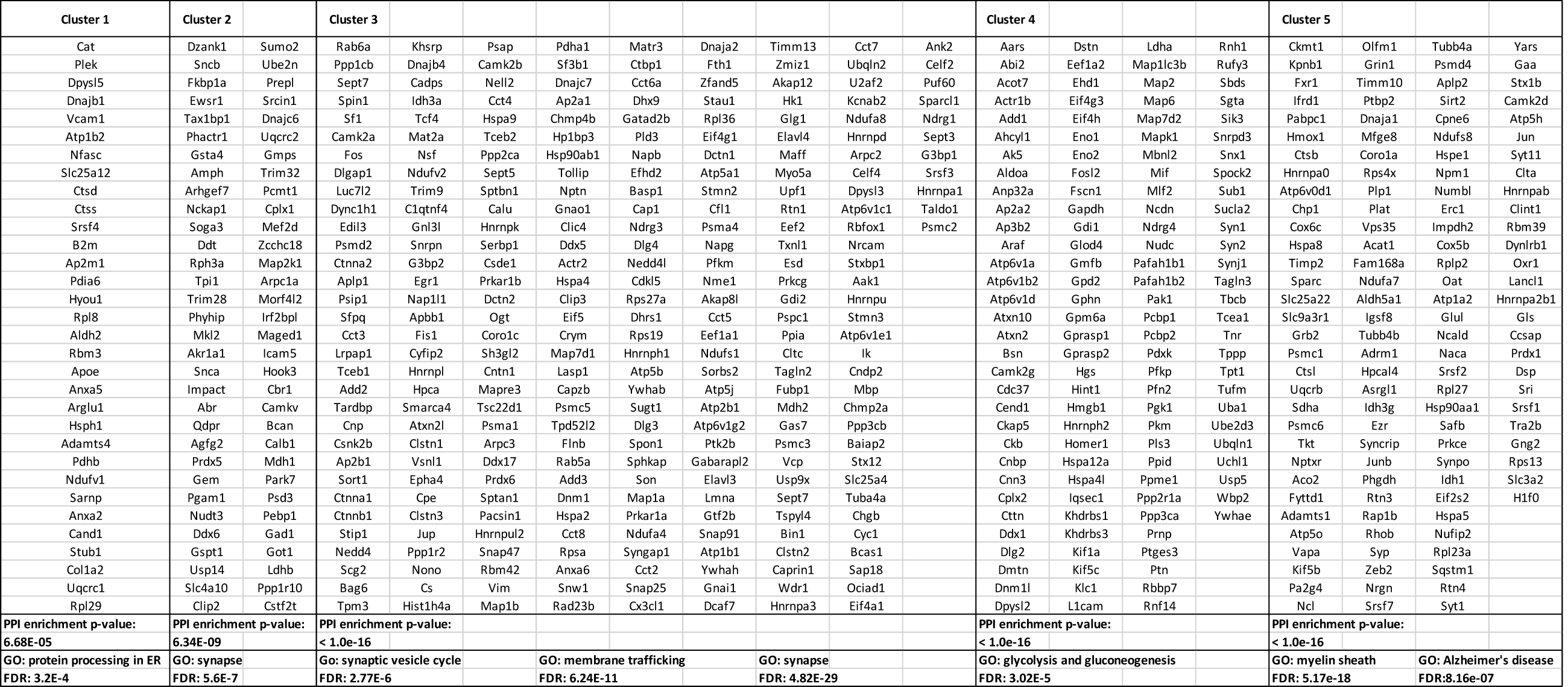
Protein identities contained within key biological pathways as detected by the hierarchical clustering depicted in **Figure 5** and **Supplementary Figures 5-10**. BONLAC-detected proteins observed in both asymptomatic and symptomatic APP/PS1 and wild-type mouse hippocampus (n = 5-7 mice/genotype/group). PPI and FDR as detected via Cytoscape and StringDb.

